# Structure-guided analysis and prediction of human E2–E3 ligase pairing specificity

**DOI:** 10.64898/2026.02.10.700855

**Authors:** Brianna Jarboe, Roland L. Dunbrack

**Affiliations:** Drexel University College of Medicine, 2900 W. Queen Lane, Philadelphia, PA 19129, USA; Institute for Cancer Research, Fox Chase Cancer Center, Philadelphia, PA, 19111 USA

**Keywords:** E2, E3, E2 enzyme, E3 ligase, AlphaFold, AlphaFold 2, AlphaFold3, machine learning, structural biology, structural bioinformatics

## Abstract

Protein ubiquitination, directed by specific E3 ligases, constitutes the primary cellular pathway for selective protein degradation. In addition to targeting proteins for degradation, ubiquitination can mediate new protein–protein interactions, and otherwise modulate protein function, thereby regulating key cellular processes such as DNA repair and immune responses. Recently, Proteolysis-Targeting Chimeras (PROTACs), and related proximity-inducing agents, have revealed the significant therapeutic potential of co-opting ubiquitin ligase activity to induce the selective degradation of disease-relevant proteins. Despite the biological and clinical significance of this pathway, fundamental gaps remain in our understanding of ubiquitination networks, particularly regarding the specificity of E2–E3 interactions and their substrate preferences. In this study, we leverage analysis of experimental structures in the Protein Data Bank (PDB) and use AlphaFold to generate structures of thousands of ubiquitin–E2–E3 ternary complexes. Using these predicted structures and complementary analyses, we develop a machine learning model to predict functional E2–E3 pairings, advancing our ability to map ubiquitination networks and providing structural insights into functional ubiquitin–E2–E3 complexes. We demonstrate the utility of our model by predicting E2 partners for 88 putative E3 ligases lacking any previously known E2 interactors. Notably, we identify a predicted pairing between UBE2C and RNF214, two proteins recently implicated in hepatocellular carcinoma separately but through interrelated pathways, suggesting a potential functional link mediated by RNF214-dependent ubiquitination in partnership with UBE2C. Additionally, we present our web-resource, UbiqCore, making the E2–E3 pairing predictions and ternary complex structures available to the scientific community (https://dunbrack.fccc.edu/ubiqcore).

**Significance Statement:** Ubiquitination is an essential protein modification that regulates nearly all aspects of biology and is mediated by dozens of ubiquitin-conjugating enzymes (E2s) and hundreds of ubiquitin ligases (E3s). These enzymes are frequently implicated in cancer and other diseases, and their activities can also be harnessed for targeted protein degradation therapies. Despite the biological and clinical significance of ubiquitination, there remains no systematic understanding of which E2s and E3s function together. Here, we combine bioinformatic analysis of experimentally determined structures and AlphaFold modeling of thousands of ubiquitin–E2–E3 complexes with machine learning to predict functional E2–E3 pairings. We also introduce UbiqCore, a web resource that makes these predictions and structures broadly accessible, providing a foundation for mapping ubiquitination networks and guiding future biological and therapeutic discovery.

## Introduction

The majority of human proteins are thought to undergo ubiquitination at some stage of their cellular lifespan, with estimates suggesting that at least 80% of the proteome is ultimately degraded via the ubiquitin–proteasome system (1–4). Protein ubiquitination is mediated by a cascade-process beginning with the ATP-dependent conjugation of ubiquitin to a conserved cysteine of an E1 enzyme (5). Ubiquitin is then transferred from the E1 to an E2 enzyme in a trans-thioesterification reaction (5, 6). Finally, an E3 ligase enables the transfer of ubiquitin from the E2 to a target substrate, typically resulting in an isopeptide bond formed between a lysine of the substrate and the C-terminus of ubiquitin (7, 8). There are two known human ubiquitin E1 enzymes, approximately 40 human E2 enzymes, and hundreds of E3 ligases thought to function in ubiquitination (9, 10). The large number of E3 ligases provide the diversity required for targeting the vast number of proteins which constitute the ubiquitome (7).

Depending on the type of E3 ligase, there are differing core domains which interact with E2 enzymes and catalyze ubiquitin transfer to substrates. In the case of RING/U-box-type E3s, either a “Really Interesting New Gene” (RING) or U-box domain protein mediates E2 binding and ubiquitin transfer. The RING protein domain is characterized by a specialized zinc-finger motif, in which two zinc ions are coordinated by cysteine and histidine residues to form a structure known as a RING finger (11, 12). While the homologous U-box domain does not bind zinc ions, alternate intra-protein interactions mediate a very similar structural motif (13, 14). Along with their closely related structures, RING and U-box E3s appear to exhibit a shared means of E2 engagement and mechanism of ubiquitin transfer (14–16). RING/U-box-type E3s, which we refer to as E3_RING_ ligases, are thought to account for the majority of human E3s (10, 17). A key feature of E3_RING_ ligase catalysis is that ubiquitin is transferred directly from the E2 enzyme to the substrate (**Fig. 1*A***). Alternately in a smaller subset of E3s, consisting of “Homologous to the E6-AP Carboxyl Terminus” (HECT) and “RING-between-RING” (RBR) E3 ligases, ubiquitin is first transferred from the E2 to a catalytic cysteine within the HECT or RBR domain of the E3 itself, via an additional trans-thioesterification, before being transferred to the substrate in a final step (18–20).

**Fig. 1.**
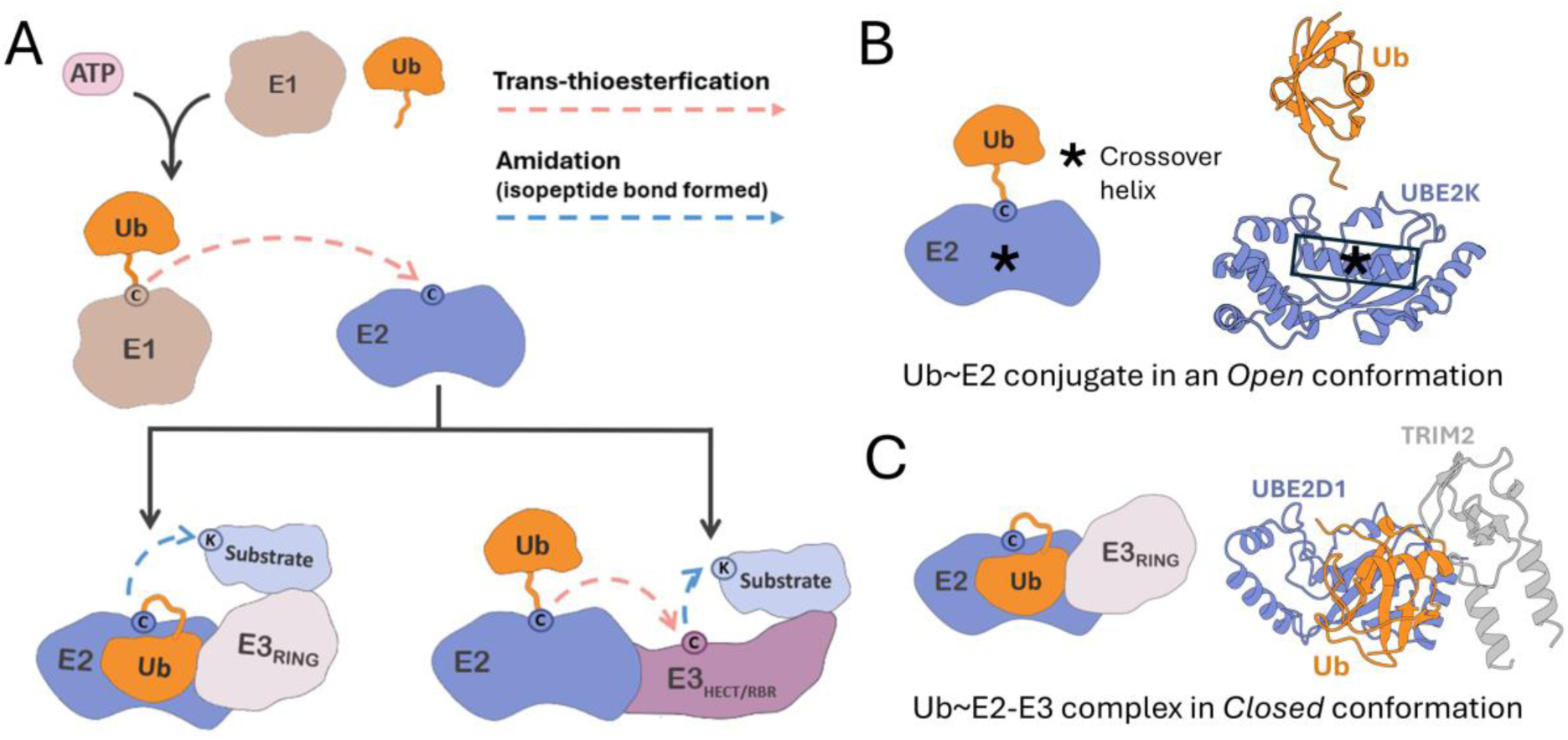
Specific mechanisms of E3_RING_ ligases. *(A)* Ubiquitin transfer mechanism of the E3_RING_ ligases versus HECT/RBR-type E3s. *(B)* Example of experimental structure of ubiquitin conjugated to E2 enzyme (Ub∼E2) in an Open conformation (PDB: 5DFL, E2 Gene Name: *UBE2K*)*. (C)* Example of experimental structure of an E3 ligase in complex with a ubiquitin*–*E2 enzyme conjugate (Ub∼E2–E3) in the Closed conformation (PDB: 7ZJ3, E2 Gene Name: *UBE2D1*, E3 Gene Name: *TRIM2*). Note: The experimentally determined structure, PDB: 7ZJ3, forms a dimer of ternary complexes but only one ternary complex is displayed for visual clarity.

Structural studies of ubiquitin∼E2 conjugates have revealed relatively free movement of ubiquitin relative to the E2, enabling both ‘Open’ and ‘Closed’ conformations (21, 22). In variants of the Open conformation, ubiquitin can make little to no contact with the E2 besides the linking bond (**Fig. 1*B***). In the Closed conformation, ubiquitin is tightly folded over the E2, such that a hydrophobic region of ubiquitin, centered about Ile44, makes significant contact with the so-called ‘crossover’ α2 helix of the E2 Ubiquitin-conjugating core (UBC) domain (9, 22). Studies investigating E3 ligases and ubiquitin∼E2–E3 ternary complex structures indicate that E3_RING_ ligases stabilize the ‘Closed’ conformation of ubiquitin∼E2 conjugates (23–25) (**Fig. 1*C***). Furthermore, evidence suggests that this may be the major means of E3_RING_ catalysis of ubiquitin transfer from E2 enzymes to substrates (26, 27). Specifically, the closed conformation is thought to better expose the ubiquitin–E2 thioester bond to nucleophilic attack by the lysine on the accepting substrate (28). This mechanism is specific to E3_RING_ ligases, as HECT and RBR-type E3s do not appear to stabilize the Closed conformation but rather enable ubiquitin transfer from variants of an Open conformation (29, 30).

One of the main outcomes of ubiquitination is the targeting of substrate proteins for proteasomal degradation. Beyond a means of merely recycling misfolded, or otherwise dysfunctional proteins, degradative regulation of protein half-life allows for precise control of protein abundance in response to changes in gene expression, making it a fundamental aspect of cellular biology (31–33). In addition to this function in global control of cellular protein levels, targeted degradation of substrates mediated by particular E3 ligases modulates the activity of major signaling pathways, such as cell cycle progression, pathogen sensing, and Wnt signaling (34–38). Besides targeting proteins for degradation, ubiquitination is a post-translational modification with key roles in DNA damage response, protein trafficking, and mediation of protein-protein interactions (39–43). Aberrant protein ubiquitination, typically caused by perturbations of E1-E2–E3 activity, underlies many human diseases including neurodegenerative conditions and numerous cancers (44–47).

Additionally, Proteolysis-Targeting Chimeras (PROTACs) and related compounds have emerged as therapeutic modalities harnessing the ubiquitin ligase pathway (48). PROTACs and similar proximity inducing agents are typically bivalent binders with one portion of the compound interacting with the protein of interest and another to an E3, resulting in drug-induced de novo ubiquitination of the target protein (49). This presents a promising avenue to target various drivers of disease for degradation, particularly so called “undruggable” proteins that are not readily targetable by conventional small molecule inhibitors (50).

Despite the fundamental importance of protein ubiquitination to cellular biology, there are still significant gaps in our knowledge of E1–E2–E3 mediated ubiquitination. This includes understanding which of the dozens of E2s and hundreds of E3s function together and which substrates are targeted by the resulting ubiquitin–E2–E3 complexes. While there have been increasing efforts to identify the substrates targeted by known E3 ligases (51–55), the identification of E2–E3 functional pairs has received much less attention. To address this gap, in this study we perform a bioinformatics analysis of experimental structures of ubiquitin–E2 and ubiquitin–E2–E3 complexes in the Protein Data Bank (PDB) and take advantage of recent major advances in protein structure prediction, specifically with AlphaFold-based tools, as well as machine learning to predict E2-E3 pairings for 88 E3 ligases without a known E2 partner.

## Results

### Distribution and specific mechanisms of human E3 ligases

Due to the sequence, structural, and mechanistic diversity among E3 ligases, a definitive list is elusive and current estimates of the total number of E3s vary. There can also be a lack of full clarity regarding which types of proteins are considered E3 ligases. While a protein containing a RING, U-box, RBR, or HECT domain—which are minimally required for direct interaction with E2 enzymes and ubiquitin transfer catalysis—is most often what is implied by the term, it is not uncommon for entire complexes, or various associated proteins (such as substrate-recruiting adapter proteins or scaffolds), to also be included in broader lists of E3 ligases (10, 56).

To get a clearer understanding of the human E3 ligase landscape, we pooled lists of established and putative human E3s from sources including UbiBrowser 2.0 (57), a web-based resource of known and predicted ubiquitin ligase/deubiquitinase–substrate interactions in eukaryotes, as well as from other studies aimed at cataloging human E3s (58, 59). We then assessed all of these for the presence of RING, U-box, HECT, and IBR domains, using a combination of Evolutionary Classification of protein Domains (ECOD) (60) and The Encyclopedia of Domains (TED) (61) domain classifications, as well as PROSITE (62) domain annotations. Proteins with HECT or U-box domains were classified as HECT and U-box E3s, respectively. All proteins containing RING domains were classified as a RING E3 if there was no IBR domain, but as an RBR E3 if the protein also contained an IBR. The remaining proteins, which lacked any of these domains, appeared primarily to be E3-complex-related proteins and/or substrate adaptors (***SI Appendix*, Table S1**). Concordant with other estimates (10, 56), our analysis showed that E3_RING_ ligases account for the vast majority of known human E3s due to the high prevalence of RING domain-containing non-RBR E3s (**Fig. 2*A***). Specifically, we identified 369 putative human E3 ligases: 318 proteins containing RING domains, 8 with U-box domains, 28 with HECT domains, and 14 with RBR domains from the culled list (one atypical E3 ligase, *G2E3*, was found to have both RING and HECT domains and was excluded from the counts of the currently recognized classes). To study a mechanistically unified class that also represents the majority of human E3s, for the remainder of this study we focus on E3_RING_ ligases.

**Fig. 2.**
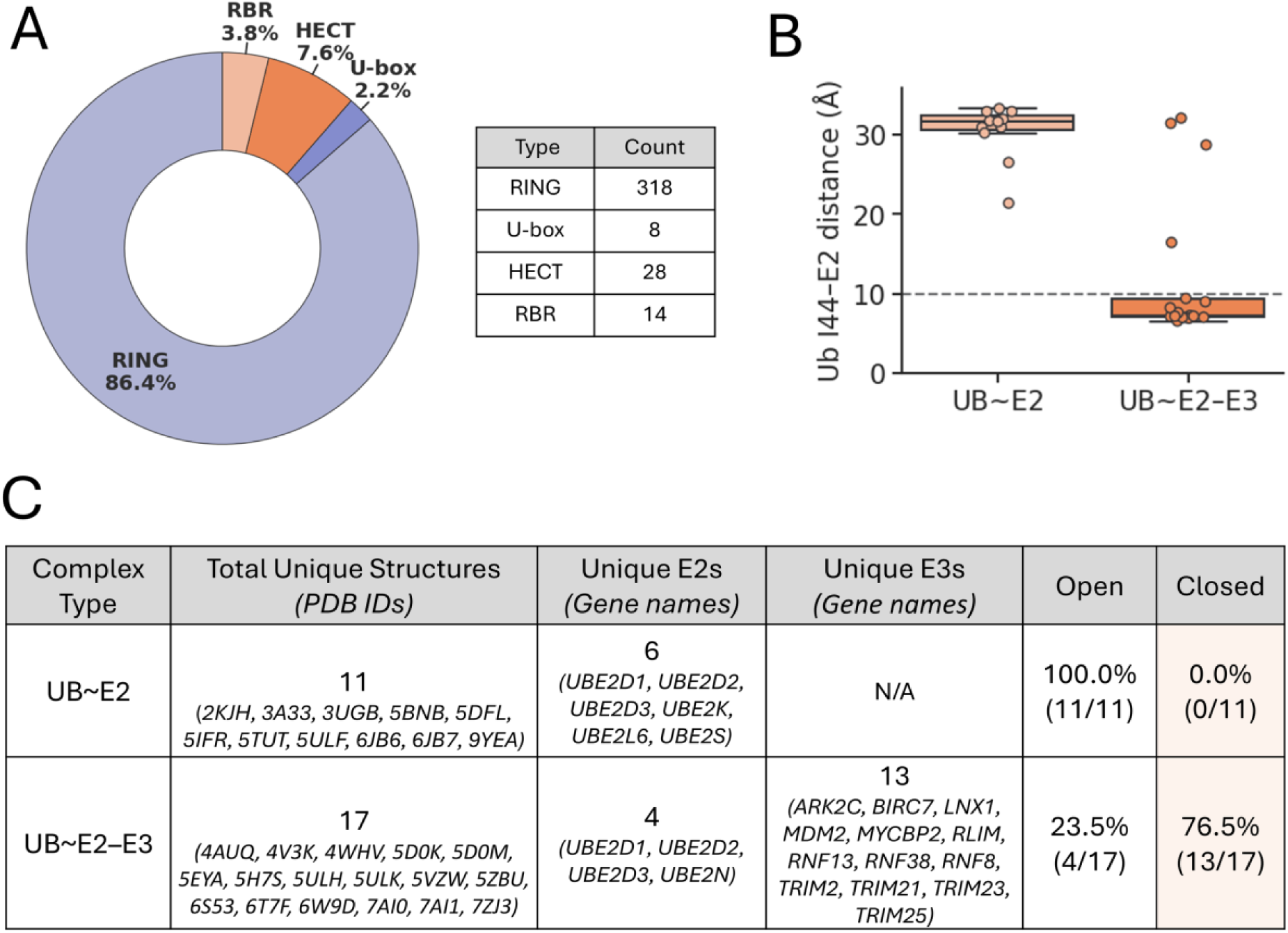
Prevalence of E3_RING_ ligases and their propensity to induce the Closed conformation of ubiquitin–E2 enzyme conjugates. (*A*) Distribution of human E3 ligases based on analysis of a compilation of putative E3 ligases from multiple sources. E3s were categorized based on the presence of HECT, RBR, U-box and RING domains. (*B*) Distances from ubiquitin Ile44 Cα to closest crossover helix Cα in ubiquitin*–*E2 enzyme conjugates alone or in complex with E3_RING_ ligases. (*C*) Table summarizing Open vs. Closed conformation analysis of ubiquitin*–*E2 enzyme conjugates alone or in complex with E3_RING_ ligases, with Open or Closed conformation designation based on 10 Å Ile44-crossover helix distance cutoff.

### Analysis of PDB structures of E2s and E3s with ubiquitin

Many studies indicate that induction of the Closed conformation of ubiquitin∼E2 conjugates likely represents a primary means of E3_RING_ ligase activation of ubiquitin transfer from E2 to substrate. To assess the prevalence of this conformation, we queried the Protein Data Bank (PDB) (63) for available experimental structures of ubiquitin–E2 conjugates alone, as well as those in complex with E3_RING_ ligases. Specifically, we considered complexes including human ubiquitin, E2 enzymes, and/or E3 ligases, and to mitigate confounding factors, excluded structures containing inhibitor or drug–like ligands, as well as complexes containing proteins other than ubiquitin, E2s or E3s. This resulted in six unique ubiquitin–E2 complexes and four ubiquitin–E2–E3 complexes in 11 and 17 PDB entries respectively.

To distinguish Closed and Open conformations in a quantitative manner, we calculated the distance between ubiquitin residue Ile44 and the E2 crossover helix across all of the available structures, revealing a clear cutoff of approximately 10 Å separating the two conformational states (**Fig. 2*B***). Notably, ubiquitin–E2 conjugates in the absence of E3 ligases exclusively adopt Open conformations, whereas more than 75% of ubiquitin–E2–E3_RING_–containing complexes are observed in the Closed conformation (**Fig. 2*C***). The small proportion of ubiquitin–E2–E3_RING_ complexes observed in the Open conformation likely reflect the intrinsic dynamic nature of ubiquitin–E2 conjugates, rather than deviations of E3 RING ligases from the Closed-conformation mechanism, as three of the four Open complexes involved the E3 ARK2C, which is known to adopt the Closed conformation during ubiquitin transfer to substrates (PDB: 5D0M) (64). Overall, these findings suggest that the Closed conformation is indeed a general structural feature of ubiquitin–E2–E3_RING_ ternary complexes.

While the experimental structures of E2s, E3s and ubiquitin–E2–E3 ternary complexes available in the PDB provide a critical basis for understanding typical E2s and E3 domains, as well as key features of functional ternary complexes, the number of these structures is relatively limited. Currently experimental structures are lacking for 5 of the 37 human E2s and 180/369 of E3s, with no experimental structures available for 164/318 of putative RING/U-box-type E3s (***SI Appendix*, Table S2**). Additionally, with only 57 ternary complex containing experimental structures in the PDB, this covers only a very small fraction of the ubiquitin–E2–E3 complexes that are likely to function in humans (***SI Appendix*, Table S3**).

### Modeling of ubiquitin, E2, and E3 containing complexes with AlphaFold

AlphaFold (65, 66) and related emerging tools present an excellent opportunity to extend our understanding of ubiquitin–E2–E3 complexes and the ubiquitination networks as a whole. To explore the potential utility of using AlphaFold (AF) in this context, we first evaluated AF3 modeling of ubiquitin–E2–E3 complexes. An important consideration was that we would not be able to include a ubiquitin–E2 thioester bond explicitly but based on the consideration that this bond serves to flexibly tether ubiquitin to an E2, we hypothesized that ternary complexes might be successfully modeled even without an explicit bond.

We first wanted to assess AF3 complex structure predictions for ubiquitin–E2–E3 complexes that were not part of its training set. To this end, we compared AF3 predicted complex structures to experimentally determined structures released to the PDB after the AF3 training dataset date cutoff of Sept. 30, 2021. We found six human ubiquitin–E2–E3_RING_ containing structures in the Closed conformation that were only released in the PDB after this date. We generated AF3 predictions for each and found that the predicted structures recapitulate expected features—such as conserved domains/motifs and proper orientation of the E3_RING_ domain at the canonical E2 binding site (**Fig. 3*A***). In addition to the ubiquitin–E2–E3_RING_ containing structures, there was also an experimental structure (PDB: 8EB0) (18) released after the AF3 training dataset cutoff containing a ubiquitin-conjugated E2 (UBE2L3) in complex with an RBR-type E3, RNF216. While RBR-type E3 ligases feature two RING domains (which flank the central IBR domain), they are known to catalyze ubiquitin ligation in a distinct manner from E3_RING_ ligases and, conversely, have been shown to stabilize an Open rather than Closed conformation (18, 67). To further evaluate AF3 predictive power in the case of a RING domain in the context of an E3_RBR_ ligase, we modeled this complex with AF3 as well. We found that AF3 did indeed recapitulate the expected Open conformation found in the experimentally determined structure.

**Fig.3.**
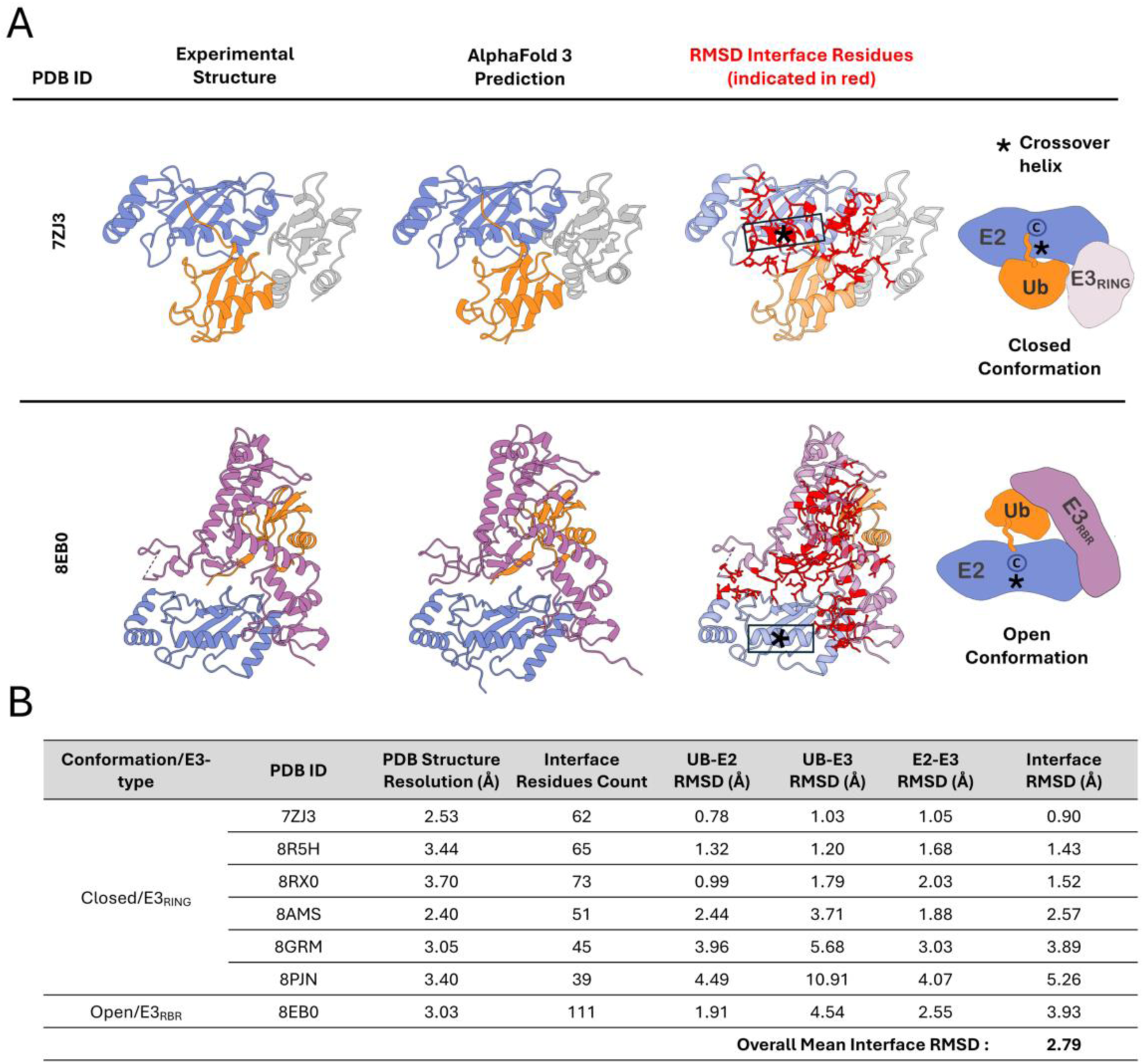
AlphaFold3 modeling of ubiquitin–E2–E3 ternary complexes. (*A*) Examples of experimental and AlphaFold3-predicted ubiquitin–E2–E3 complex structures. Complexes are viewed ‘overhead’ with respect to ubiquitin tail conjugation to E2 for visualization of interfaces. For each complex, the experimental structure (left), AlphaFold3 prediction (middle), and superimposed interface residues used for RMSD calculation (right, in red) are shown. An example of a ubiquitin–E2–E3_RING_ complex in the Closed conformation (top row) and a ubiquitin–E2–E3_RBR_ complex in an Open conformation (bottom row) are shown. PDB:7ZJ3 contains E2, UBE2D1 and E3, TRIM2. PDB: 8EB0 contains E2, UBE2L3 and E3, RNF216. (*B*) Table of interface RMSD values calculated by comparing experimentally determined structures of ubiquitin–E2–E3 complexes (PDB IDs listed) with their corresponding AF3-predicted models. The table includes the resolution of the experimental structures, and the number of interface residues used in the RMSD calculations, defined as residues with Cα atoms within 8 Å across inter-protein interfaces of the complex. All experimental structures were released after AF3 training dataset cutoff of Sept. 30, 2021.

To quantitatively assess the relative positioning of ubiquitin, E2, and E3 components within the complexes, as well as the overall quality of interface modeling, we calculated interface RMSD values between experimental structures and AF3-predicted complexes. For the majority of ubiquitin–E2–E3_RING_ complexes, the interface RMSD was below 3 Å, with an overall mean of 2.79 Å, indicating good general agreement between predicted and experimental structures (**Fig. 3*B***). A subset of structures, however, exhibited higher RMSD values, most notably PDB: 8PJN with an interface RMSD of 5.26 Å. An important consideration is that, due to the limited number of structures published after the AF3 training data cutoff, we included experimental complex structures containing additional protein components (beyond ubiquitin, E2, and E3), as well as structures harboring post-translational modifications such as phosphorylation. In both experimental structures, 8PJN and 8GRM, the E3s of the ternary complex form heterodimers with a partner E3 (although only one E3 makes primary contact with the E2), so we additionally generated models including the E3 heterodimer partners for these structures (***SI Appendix,* Fig. S1*A***). When the partner E3 was included in the AF3 model for 8PJN, the interface RMSD improved markedly, decreasing to 1.43 Å (***SI Appendix,* Fig. S1*A***), indicating that the lack of the dimer partner was the primary source of the observed discrepancy for this structure. We also modeled additional ubiquitin–E2–E3_RING_ complexes not found in the PDB, including examples with putative E2–E3_RING_ functional pairs, and E2–E3_RING_ pairs with no evidence of interaction, and found that complexes could be modeled in both Open or Closed conformations depending on the E2–E3_RING_ pair (***SI Appendix,* Fig. S1 *B* and *C***). Overall, we found that despite the absence of an explicit thioester bond, AF was able to model ubiquitin–E2–E3 ternary complexes, producing plausible structures with expected key features, with the potential of both the Closed and Open conformations, dependent on the type of E3 ligase or the particular combination of E2 and E3_RING_.

### Modeling ubiquitin–E2–E3 ternary complexes of E2–E3 pairs and non-pairs

Intrigued by the presence of both Closed and Open conformations among the AF3 ternary complex predicted structures, we next evaluated whether prediction of a Closed vs Open conformation might correlate with ubiquitin–E2–E3_RING_ complexes of functional E2–E3_RING_ pairs vs non-pairs. Wet lab screens for likely E2–E3 pairs have been performed using various protein-protein interaction (PPI) detection assays, but to generate as comprehensive a set of E2–E3_RING_ pairs as possible, as inferred by detected PPI, we used the of protein-protein interaction database STRING (68), which we found to encompass all major sources of PPI data. For an input list of E2s with which to query STRING, we used the *Ubiquitin conjugating enzymes* E2 HGNC gene group (69), filtering out predicted pseudogenes (***SI Appendix*, Table S4**). We also removed *AKTIP*, which encodes a related but non-E2 protein, and *BIRC6*, which encodes a very large atypical E2/E3 combination protein. As for E3s, our focus in this study is RING/U-box-type E3 ligases, which comprise the vast majority of E3s, and operate via a shared mechanism of ubiquitin transfer involving induction of the Closed conformation of the ubiquitin–E2 conjugate. However, for a comprehensive assessment of all E3s bearing RING domains, RBR E3s were also included initially in our analysis.

To establish a ‘pair-enriched’ set, we queried the STRING database for E2 and E3 physical interactions and collected any intersecting pairs. From this pairs list, we included only interactions with a STRING experimental data score of medium or higher (which corresponds to a numerical score of ≥ 400). This resulted in the identification of 402 E2–E3 pairs (**Fig. 4*A***), comprising 31 unique E2s and 146 unique E3s. For a control, or ‘non-pair enriched’ set, we generated all possible 4,526 E2-E3 pairs from the unique E2s and E3s from the pair-enriched set. From this list, after we filtered out all pairs with *any* level of evidence for interaction in the STRING database this left 3,463 pairs, from which we made a final random selection of 402 non-pairs to match the number of the pair enriched set (**Fig. 4*A***).

**Fig. 4.**
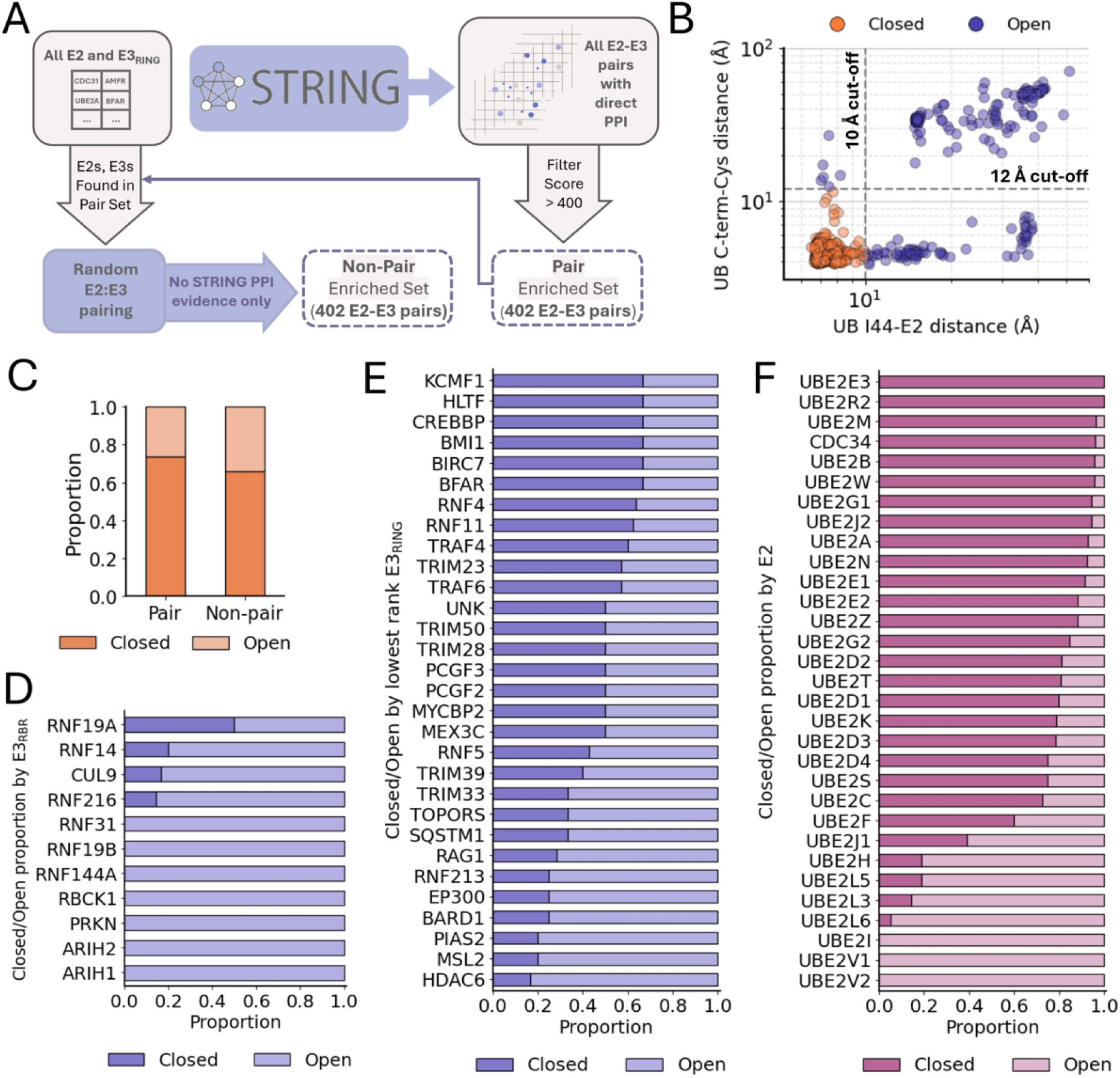
Modeling of ubiquitin–E2–E3 ternary complexes of E2–E3 pair and non-pair sets with analysis of Closed versus Open conformations. (*A*) Schematic of E2–E3 pair vs non-pair set generation using STRING database. (*B*) Scatter plot of ubiquitin Ile44 to E2 crossover helix distance vs ubiquitin Gly C-terminus to E2 catalytic cysteine distance for all structures. (Note: Structures containing the E2s UBE2V1 or UBE2V2, which lack a catalytic cysteine, are not shown in this plot. However, all such structures exceeded the 10 Å Ile44–E2 distance cutoff and were assigned Open conformation designation.) (*C*) Proportion of ternary complexes in a Closed vs Open (non-Closed) conformation for pair and non-pair set. Closed/Open proportion by E2 (*F*), E3_RBR_ (RBR-type E3) (*D*), and the 30 lowest Closed proportion ranking E3_RING_ (RING/U-box-type E3) (*E*) (E2s and E3s referred to by gene name).

Next, we generated AF3 models for all of the E2–E3 pairs in complex with ubiquitin. For each complex, we generated 50 models, using 10 random seeds and 5 diffusion samples per seed, then selected the top-ranked model among them. We then determined whether each complex was in the Closed conformation or an alternate conformation, all termed Open. As noted, the primary determinant of the Closed conformation is contact between the region of ubiquitin centered around Ile44 and the crossover helix of the E2. In the correct orientation of ubiquitin relative to the E2, the C-terminal tail of ubiquitin is positioned near the E2’s catalytic cysteine. To establish quantitative thresholds for identifying the Closed conformation, we plotted the distances between ubiquitin Ile44 and the E2 crossover helix versus the distances between the ubiquitin C-terminal glycine and the E2 catalytic cysteine, across all complexes (**Fig. 4*B***). Based on these distributions, we established cutoffs of 10 Å and 12 Å for the distances, respectively.

While there was a slightly greater propensity of the pairs vs non-pairs to form the Closed conformation, most of the complexes were modeled in the Closed conformation in both cases (**Fig. 4*C***). True to our understanding of their physiological functioning, complexes of E3_RBR_ had an increased propensity to form an Open conformation compared to complexes of E3_RING_ ligases. Analysis of the proportion of the Closed/Open complex conformation by E3 showed that for most E3_RBR_ all complexes were in the Open conformation (**Fig. 4*D***), while even the lowest ranked E3_RING_ had some proportion of complexes in the Closed conformation (**Fig. 4*E***). This is consistent with previous studies indicating that the E2-binding RING domain of RBR E3s paradoxically stabilizes the open conformation of ubiquitin–E2 conjugates (18, 29, 67, 70). This highlights the fact that RBR E3s operate via mechanisms fundamentally distinct from E3_RING_ ligases—despite their shared inclusion of RING domains—and are therefore best considered separately in such contexts.

When we examined Closed/Open conformation proportion by E2, we also found physiologically relevant correlates. The ubiquitin-conjugating enzyme E2 variants 1 and 2 (UBE2V1 and UBE2V2) (71, 72) both lack the conserved cysteine needed for ubiquitin conjugation, and are therefore catalytically dead E2s. All models with these E2s were found to be in an Open conformation (**Fig. 4*F***). Additionally, all complexes with the E2 enzyme SUMO-conjugating enzyme UBC9 (UBE2I), which is known to function with the ubiquitin-like protein, SUMO, rather than ubiquitin (73), were also all in Open conformations (**Fig. 4*F***). There was also a very low proportion of complexes in the Closed conformation for E2s Ubiquitin-conjugating enzymes UBE2L3 and UBE2L6, which previous work indicates are likely E2 enzymes that are specific to HECT/RBR E3 ligases (74–76). Based on these results for the remaining analyses, we exclude all RBR E3s and the HECT/RBR specific E2s UBE2L3, UBE2L6 (as well as UBE2L5, which is closely related to UBE2L3/6 and a potential pseudogene). Additionally, we exclude the catalytically inactive E2s UBE2V1, UBE2V2, as well as the SUMO-specific E2, UBE2I, in all the proceeding analyses.

### Analysis of predicted ternary complex confidence metrics

Notwithstanding the noted exceptions, which compellingly align closely with specialized E2/E3 functionality, the majority of complexes were modeled in the Closed conformation, regardless of whether they belonged to the pair or non-pair set. As such, we hypothesized that AF3 confidence scores for Closed-conformation complex structures might correlate with pair vs non-pair status. Indeed, based on the interface predicted template modeling (ipTM) scores produced by AF3 and interface-averaged predicted aligned error (PAE) values, there was significantly higher confidence in complex interfaces from the pair compared with the non-pair sets (**Fig. 5 *A* and *B***). Notably, the median interface-averaged PAE across true pair complexes was < 3.15 Å for all three interfaces (UB–E2, UB–E3, and E2–E3), whereas non-pair complexes showed median interface-averaged PAE values ranging from 4.71 to 5.91 Å. We similarly evaluated interaction prediction score from aligned errors (ipSAE) (77) and Predicted DockQ version 2 (pDockQ2) (78) scores, which both were significantly more favorable in the case of complexes of the pair vs non-pair sets (**Fig. 5 C and D**). In additional analyses of the ternary complex models of our pair and non-pair sets, we used the Rosetta InterfaceAnalyzer (79) application to assess complex interface metrics, including binding energies and hydrogen bonding. We found that there were a lower number of unsatisfied interfacial hydrogen bonds (**Fig. 5*E***) and more favorable binding energies (**Fig. 5*F***) in pair vs. non-pair complexes, particularly for the ubiquitin–E2 interface.

**Fig. 5.**
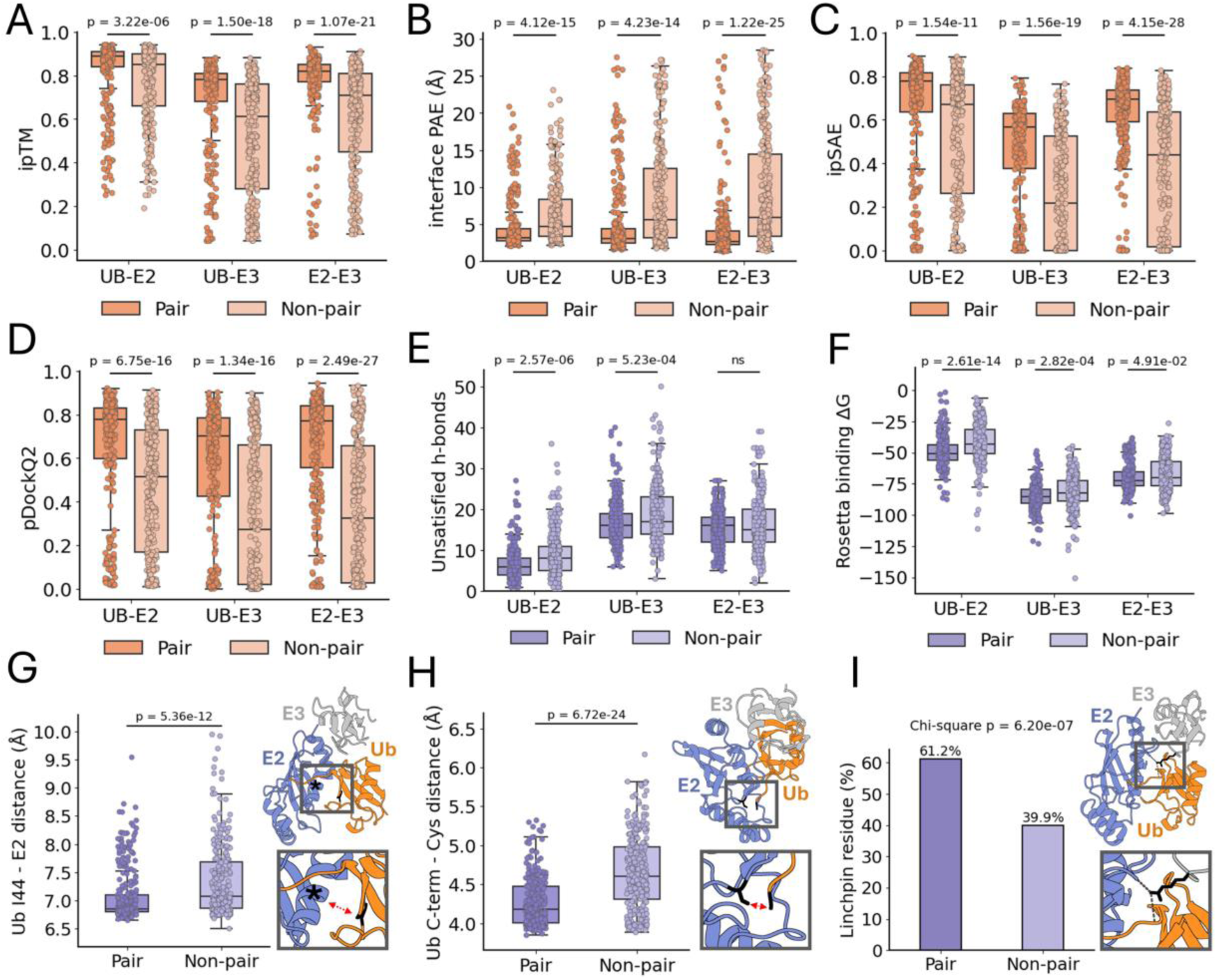
Interface confidence metrics, energetics, and structural features of pair and non-pair ubiquitin–E2–E3 Closed conformation ternary complex structures. Complex interface confidence metrics ipTM (*A*), average interface PAE values (based on 8 Å Cα-Cα interface cutoff) (*B*), ipSAE (*C*), and pDockq2 (*D*) plotted for each complex interface. Number of unsatisfied hydrogen bonds at interfaces (*E*) and complex interface binding energies (Rosetta units) (*F*), both determined using Rosetta InterfaceAnalyzer. (*G*) Distances from ubiquitin Ile44 Cα to closest crossover helix Cα. Crossover helix marked by asterisk (*) in example structure, Ile44 colored in black. (*H*) Distances from ubiquitin terminal glycine Cα to the E2 active-site cysteine carbonyl carbon, stick structures of both residues are indicated in black in example structure. (*I*) Proportion of complexes featuring linchpin residue in pair and non-pair complexes. Arginine linchpin residue indicated in black in example structure. Two-sided Mann–Whitney U tests were used to calculate p-values unless otherwise noted; ns denotes p > 0.05. For panels (*A* and *C-I*), *n* = 294 (pair) and *n* = 273 (non-pair). For (*B*), group sizes vary by interface (due to exclusion of structures lacking interface residues within the 8 Å cutoff): UB–E2: *n* = 294 (pair), *n* = 273 (non-pair); UB–E3: *n* = 290 (pair), *n* = 249 (non-pair); E2–E3: *n* = 292 (pair), *n* = 273 (non-pair).

### Analysis of ternary complex structural features

To further assess the ternary complexes of pairs versus non-pairs, we examined specific structural features known to be important in active ternary complexes. As discussed, the closed conformation is a critical mechanism by which E3_RINGs_ activate ubiquitin–E2 conjugates for ubiquitin transfer to substrates. A key aspect of this Closed conformation is the interaction between a hydrophobic region of ubiquitin (centered around Ile44) and the E2 crossover helix. Up to this point, we have classified ternary complex conformations as either Closed or Open in a binary manner. To introduce more gradation into this analysis, we compared the specific distance between ubiquitin Ile44 and the E2 α2-crossover helix in the structures. We found that there was a modest but statistically significant (p = 5.36e-12) decrease in the mean distance in pair (median = 6.86 Å) vs non-pair complexes (median = 7.08 Å) (**Fig. 5*G***). In addition to the general positioning of ubiquitin in the Closed conformation, the orientation of the ubiquitin C-terminal tail (residues 71–76) relative to the E2 has also been identified as an important feature of the active ternary complex (9, 80, 81). As a quantitative measure of this positioning, we determined the distance between the ubiquitin C-terminus and the E2 active site cysteine. For this distance we similarly found a limited but statistically significant (p = 6.72e-24) decrease in the mean distance in pair (median = 4.18 Å) vs non-pair complexes (median = 4.60 Å) (**Fig. 5*H***).

Another feature that has been found in functional ternary complex structures is a so-called ‘linchpin’ residue within the E3 ligase that mediates contact with both the E2, and the C-terminal tail of ubiquitin. Though not required in all cases, this residue is frequently present in functional ubiquitin–E2–E3_RING_ complexes (82–87). This linchpin residue has been found to be located within the finger motifs of E3 ligases and is often an arginine residue, although other amino acids are possible (84, 85). To identify any potential linchpin residues, we analyzed ternary complex structures for any E3 residues with hydrogen bonds to both the ubiquitin tail and the E2. We found that all structures had at most one such bridging residue within the E2 binding domain of E3s, and that it was an arginine residue in the vast majority of cases, which was consistent with what we found in experimental ubiquitin∼E2–E3_RING_ ternary complexes (***SI Appendix,* Fig. S2 *B* and *C***). The occurrence of an E3 bridging residue was more common in pair (61.2%) vs non-pair (39.9%) complexes (**Fig. 5*I***), the proportion of pair complexes featuring linchpin residues aligned closely with experimental ubiquitin∼E2–E3_RING_ ternary complexes (**Fig. S2*A***).

### Machine learning model for E2–E3 pair prediction

With significantly more favorable confidence metrics, binding energies, and other structural aspects correlated with the pair vs non-pair AF3 structures, we next investigated the efficacy of leveraging these features in a machine learning model to predict pair vs non-pair E2–E3s. To this end we trained an XGBoost (88) classifier model with 80% of the data set, with 20% reserved for validation (**Fig. 6*A***). After feature optimization and tuning we achieved a mean receiver-operator characteristic area-under-the-curve (ROC-AUC) value of 0.861 and an overall accuracy of 79.2% with 5-fold nested cross-validation (**Fig. *6 B* and *C***).

**Fig. 6.**
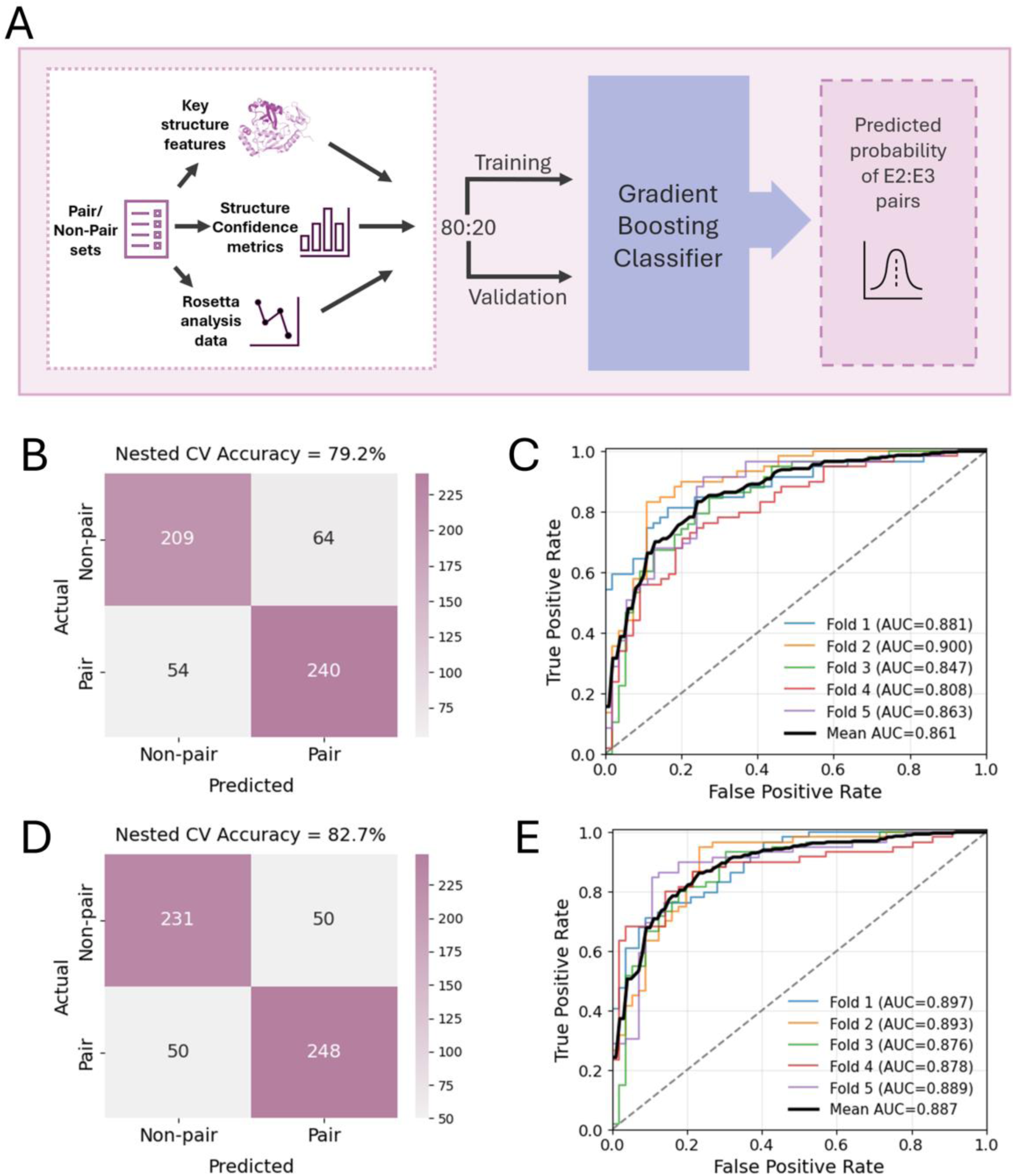
Machine learning model for predicting E2–E3 pairs from structural features and confidence metrics of ubiquitin–E2–E3 ternary complexes. (*A*) Schematic of classifier machine learning model input and training-validation split for nested cross-validation. Confusion matrix (*B*) and receiver operating characteristic (ROC) curves (*C*) for AF3 structure-based classifier resulting from 5-fold nested cross-validation. Confusion matrix (*D*) and receiver operating characteristic (ROC) curves (*E*) for CF structure-based classifier resulting from 5-fold nested cross-validation.

In addition to AF3, we wanted to evaluate the performance of AF2 predicted ternary complex structures. Specifically, we employed the ColabFold (89) (CF) implementation of AF2 for complex modeling. Generally, our analysis of CF ternary complex confidence metrics, Rosetta interface values, and other structural features revealed patterns broadly consistent with those observed in AF3 complex structures (***SI Appendix*, Fig. S2 *D–G*, Fig. S3, Fig. S4**). Our XGBoost classifier model trained from CF ternary complex data yielded an improved mean ROC-AUC value of 0.887 and an overall accuracy of 82.7% with 5-fold nested cross-validation (**Fig. *6 D* and *E***).

Based on the improved cross-validation performance of our classifier trained on CF complex features, we trained a final model using the full CF dataset and applied it to predict E2 partners for all E3_RING_ ligases lacking known E2 interactors. Specifically, we included all putative E3_RING_ ligases with no physical interaction with any E2 in the STRING database (at any level of evidence), resulting in a set of 88 E3s. Each of these was modeled in ternary ubiquitin–E2–E3 complexes with all E2 enzymes. The resulting predicted structures were analyzed, then classified as either pair or non-pair using our trained model, with the predicted E2 partners for each E3 ligase reported in ***SI Appendix*, Data S3**. From these predictions, we identified ten known/putative E3 ligases with a uniquely predicted E2 partner (**Fig. 7*A***).

**Fig. 7.**
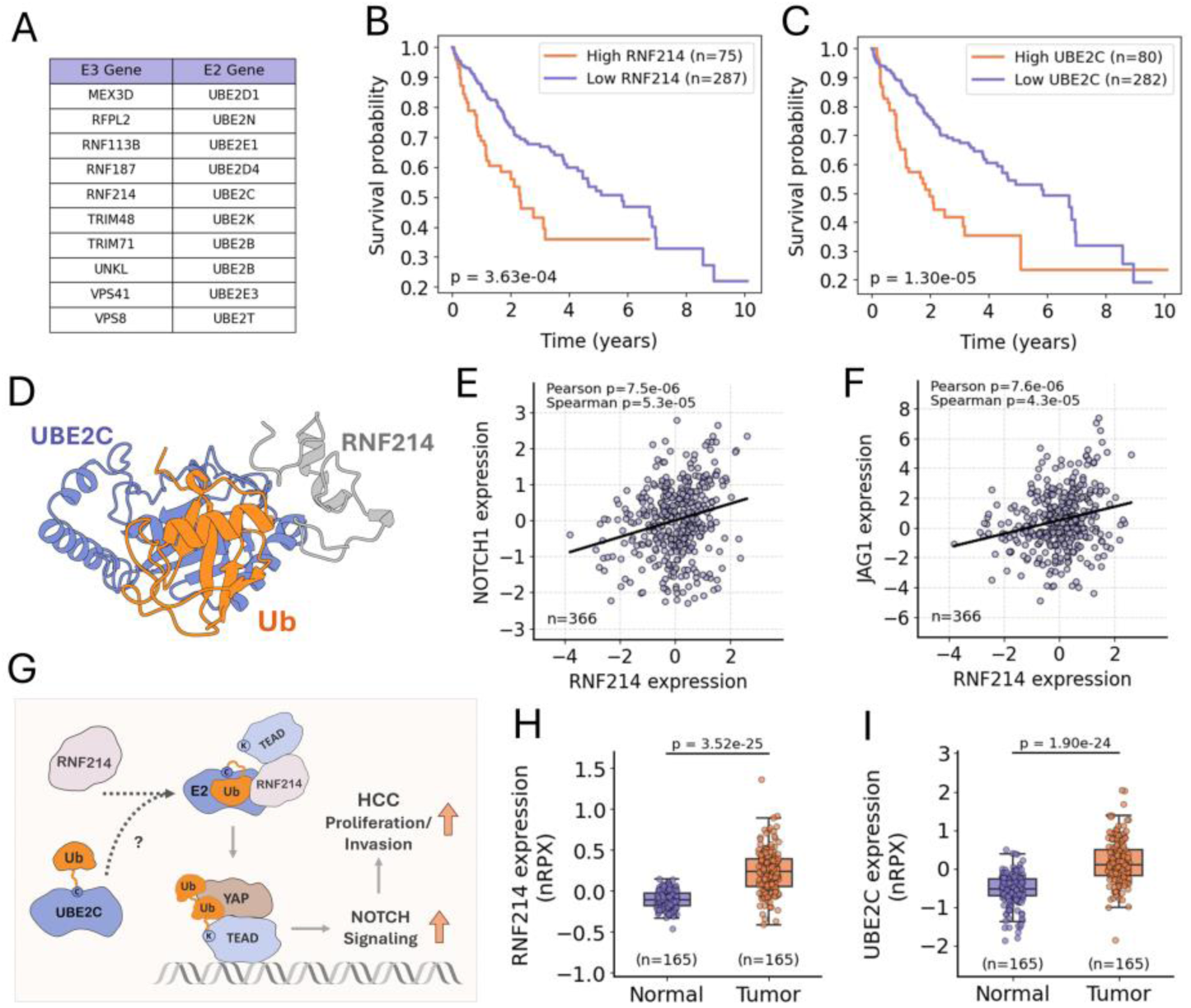
Predicted E2–E3 pairing of UBE2C and RNF214 and their convergent involvement in HCC pathology. (*A*) Table of E2 enzyme pairs for each of the putative E3 ligases with a single predicted E2 pair (Note: To our knowledge ubiquitin ligase activity has not been established for RFPL2, RNF113B, UNKL, VPS41, or VPS8). Kaplan-Meier survival plots for HCC patients stratified by tumor expression of RNF214 (*B*) or UBE2C (*C*), p-values calculated using log-rank test. Patients were divided into ‘High’ and ‘Low’ expression groups based on a best expression FPKM cutoff for each gene (RNF214: 6.18 FPKM; UBE2C: 47.29 FPKM). The Cancer Genome Atlas (TCGA) data obtained from the Human Protein Atlas (97) (www.proteinatlas.org). (*D*) Ubiquitin-UBE2C-RNF214 ternary complex ColabFold (AF2). Scatter plots of RNF214 mRNA expression z-scores vs NOTCH1 (*E*) and JAG1 (*F*) mRNA expression z-scores in HCC patient samples. Each point represents an individual tumor sample, data were obtained from cBioPortal (98) (www.cbioportal.org) (Study: TCGA PanCancer Atlas). (*G*) Schematic of the predicted UBE2C–RNF214 pairing, integrating previous findings implicating RNF214 in the activating ubiquitination of TEAD and UBE2C in the upregulation of Notch signaling, both within the context of HCC. Boxplots showing RNF214 (*H*) and UBE2C (*I*) normalized relative protein expression (nRPX) values from global protein abundance mass spectrometry measurements in tumor and adjacent normal liver tissue from HCC patient samples. P-values were calculated using the Wilcoxon signed-rank test. Clinical Proteomic Tumor Analysis Consortium (CPTAC) data obtained from the Human Protein Atlas (www.proteinatlas.org).

Among these, RNF214 stood out as a particularly compelling candidate due to its recently uncovered role in promoting hepatocellular carcinoma (HCC). RNF214 is highly expressed in HCC and is associated with poor patient prognosis (90, 91) (**Fig. 7*B***). Recent studies have demonstrated that RNF214 enhances tumor cell proliferation and invasion via activation of the YAP–TEAD signaling axis in HCC (91). Specifically, RNF214 mediates non-degradative ubiquitination of TEAD transcription factors, which enhances their interaction with YAP to increase YAP–TEAD transcriptional activity (91). However, the E2 enzyme partnering with RNF214 in this ubiquitination, or more broadly in its activity generally, has not been identified.

Our model predicted UBE2C as a unique E2 partner for RNF214 (**Fig. 7*A***) – the corresponding predicted ubiquitin–UBE2C–RNF214 ternary complex is shown in **Fig. 7*D***. Notably, UBE2C is also associated with poor prognosis in HCC (**Fig. 7*C***). Like RNF214, UBE2C has also been recently shown to promote HCC cell proliferation and invasion, in the case of UBE2C, this activity was specifically linked to increased Notch signaling but the specific mechanism was not elucidated (92). Importantly, interaction between YAP–TEAD and Notch pathways is well established, and both NOTCH1 and JAG1 (encoding a Notch receptor and ligand, respectively) have been identified as transcriptional targets of the YAP–TEAD protein complex (93–96). Furthermore, in HCC patient samples, RNF214 expression positively correlated with both NOTCH1 and JAG1 expression (**Fig. 7 *E* and *F***). Together, these findings, along with our model’s predicted RNF214–UBE2C pairing, support a convergent mechanism in which UBE2C and RNF214 may function as a E2–E3 pair to enhance YAP–TEAD transcriptional activity through TEAD ubiquitination, leading to upregulation of Notch signaling and promotion of HCC progression (**Fig. 7*G***). Additionally, both UBE2C and RNF214 were found to be significantly upregulated at the protein level in tumor tissue compared to adjacent normal tissue in HCC patients (**Fig. 7 *H* and *I***). This suggests that, in addition to representing potential therapeutic targets, UBE2C and RNF214 may be promising PROTAC candidates with enhanced tumor-specific activity.

### Ubiquitin Conformational Resource (UbiqCore)

To make the results of this work readily accessible to the scientific community, we developed the web resource UbiqCore (https://dunbrack.fccc.edu/ubiqcore). This website provides the ubiquitin–E2–E3 ternary complex structures generated in this study for user download and visualization (**Fig. 8**), together with the associated data and functional pair predictions.

**Fig. 8.**
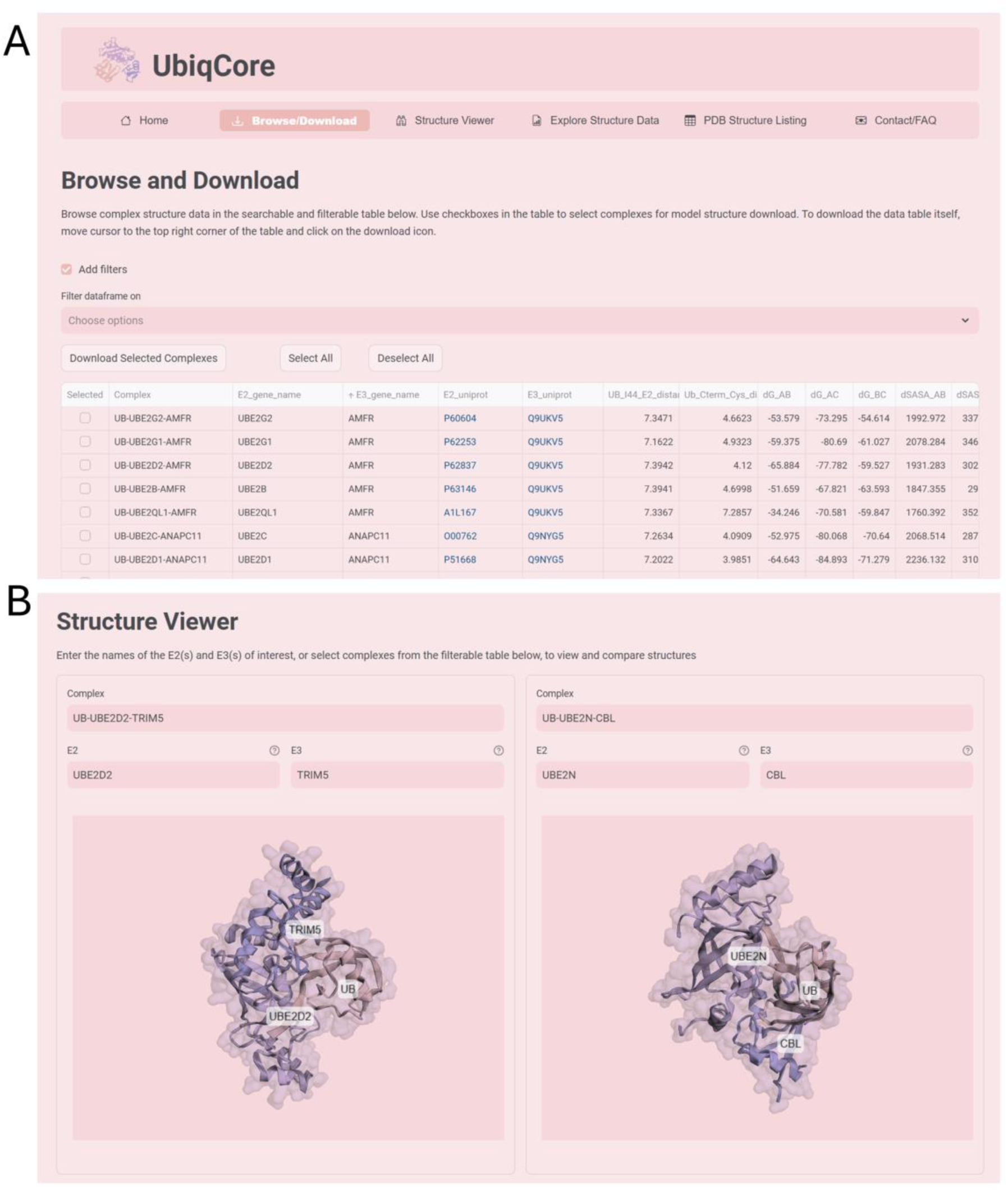
UbiqCore web resource. Representative screenshots of the UbiqCore website showing the structure download (*A*) and visualization (*B*) pages.

## Discussion

The notable gap in knowledge of functional E2–E3 pairs motivated us to perform a structural bioinformatics analysis of ubiquitin–E2–E3 experimental structures in the PDB and to explore AlphaFold predictions of ubiquitin–E2–E3 complex structures. We found that most ternary ubiquitin-E2-E3_RING_ experimental structures were in a closed conformation, while most binary ubiquitin-E2 structure were open in an open state. We also found that AF2 and AF3 are capable of predicting rational structures for ubiquitin–E2–E3 ternary complexes, with various conformations of ubiquitin positioning (i.e. Closed and Open conformations). We then evaluated complex interfaces confidence metrics, binding energetics, and other intermolecular interactions in E2–E3 pair vs non-pair sets and found that E2-E3 pairs generally scored better on all metrics than non-pairs.

In turn, we combined these results as input to an XGBoost classifying model to identify E2–E3 pair vs non-pairs. While our training-validation dataset was constructed based on PPI data (rather than direct evidence of ubiquitin ligation activity), the pair set would be expected to be highly enriched in functional pairs relative to the non-pair set. Still, there remains the possibility of a nontrivial number of functional pair false-positives (e.g. true PPIs but not functional E2–E3 pairs capable of ubiquitinating substrates), as well as non-pair false-negatives (e.g. E2–E3s which function together but for which there is no existing experimental data). Despite the introduction of such noise, robust machine learning algorithms like XGBoost can uncover key underlying patterns that distinguish true positives from negatives, even in the case of imperfect training datasets (97), as is evident in this application, based on strong model performance in cross-validation.

As a demonstration of our model’s utility, we predicted E2 pairs for 88 E3 ligases lacking any previously established E2 enzyme pairs. From the resulting set of predicted pairings, we identified UBE2C as a unique predicted E2 partner for the cancer-associated E3 ligase RNF214, which lacks any known E2 interactors. While wet lab validation is beyond the scope of this study, this prediction was supported by pathway context and expression correlations in HCC and highlights the potential of our model to generate targeted hypotheses for future experimental investigation.

Considering the fundamental nature of ubiquitination in virtually all cellular processes, the relatively limited knowledge of functional E2–E3 pairs is a significant unmet need. Furthermore, PROTACs and related compounds are exemplary of the therapeutic potential of harnessing this pathway, whether in hijacking ubiquitin–E2–E3 complex activity to induce ubiquitination of de novo targets, or to otherwise modulate/disrupt ubiquitination. The results of this work, made readily accessible through our website UbiqCore, advance efforts toward a comprehensive mapping of functional E2–E3 pairs. Calculations of all possible E2–E3 pairs are ongoing and will be added to the website in due course. The resulting E2–E3 pair predictions and structural insights provide a framework for future investigations into ubiquitination networks and the therapeutic targeting of the ubiquitin system.

## Methods

### Identifying E2s and E3s for analysis

A listing of human E2 genes was obtained from the HGNC website (url: https://www.genenames.org/data/genegroup/#!/group/102), under gene group ‘Ubiquitin conjugating enzymes E2 (UBE2)’. For the purposes of this work, all genes listed as pseudogenes were filtered out. Additionally, *AKTIP* (which encodes a non-E2 protein) and *BIRC6* (which encodes a very large atypical E2/E3 hybrid) were also excluded from the list for these analyses (Note: *BIRC6* was included in PDB queries for general determination of E2/E3 containing experimental structures). An initial listing of known/putative list of E3s was culled from multiple sources (57–59). Proteins listed as substrate adapters or other related E3 complex components without catalytic domains were excluded. UniProt IDs were mapped for all remaining proteins, and all unique proteins were then analyzed for the presence of RING, U-box, IBR, and HECT domains. Final lists of E2 and E3 UniProt IDs are provided in ***SI Appendix*, Data S1**.

### E3 ligase domain identification

Both ECOD (60) and TED (61) domain classifications, as well as PROSITE domain annotations (retrieved from UniProt) were used to identify RING, U-box, IBR, and HECT domains in our culled list of E3s. For E3-type classification, any ECOD RING/U-box, HECT, and IBR domains for each protein were identified by keyword matching domain ‘T-names’. Similarly, UniProt PROSITE domain annotations for each protein were retrieved from https://uniprot.org, and RING/U-box, HECT, and IBR domains were identified by keyword matching. Any proteins with a HECT domain were flagged as a HECT-type E3, and those with U-box domains as U-box-type. Proteins containing RING domains, as well as IBR domains, were flagged as RBR-type, and those with RING domains (but lacking IBR domains) as RING-type E3s. For AF3 and ColabFold (CF) complex modeling only RING or U-box domain containing portions of E3 protein sequences were used. To determine these RING/U-box domain containing truncated protein portions, ECOD and TED domains were prioritized since these are structure-based annotations.

For each protein identified to have one or more ECOD RING/U-box domain(s), the start and end of the domains were used for the truncation. If overlap was found with a TED domain, the start and end was expanded to cover both ECOD and TED domains. By including E3 protein regions encompassing all RING/U-box domain(s), non-RING/U-box portions between these domains are necessarily included for the E3 ligases containing more than one RING/U-box domain. Additionally, in cases of ECOD domain overlap with discontinuous TED domains, non–RING/U-box regions of E3 proteins are also included. Some constructs we used are therefore much longer than a single RING/U-box domain. However, we found that AlphaFold could still model the expected interactions of the RING/U-box domain with E2s and ubiquitin, even in the presence of extra sequence/domains. For cases where there was no RING/U-box ECOD domain, but a RING or U-box domain was identified from UniProt PROSITE annotation, the start and end of the PROSITE RING/U-box domain(s) were used to determine the truncated protein portion.

### PDB E2 and E3 containing structure identification

We queried the RCSB Protein Data Bank (PDB) (63) for all entries including UniProt IDs/Gene names of any E2 and E3 list members. The entries were filtered for human proteins/complexes containing any relevant E2 or E3. For comparative Closed/Open conformation analysis of experimental ubiquitin∼E2 conjugates and ubiquitin∼E2–E3_RING_ complexes containing structures, only those in which ubiquitin was recorded to have be conjugated to the E2 enzyme were used, structures where such a linkage could not be confirmed from the PDB entry or associated documentation were excluded. Additionally, structures that had small molecule inhibitor/drug like molecules or additional proteins other than the ubiquitin, E2, and E3 of the ternary complex were excluded from this analysis.

### Structure retrieval and visualization

All experimental structures were retrieved from the RCSB PDB. Both experimental and predicted structures were visualized using UCSF ChimeraX version 1.10.1 (100). Full length PDB codes for all structures referenced are as follows, pdb_00002kjh, pdb_00003a33, pdb_00003ugb, pdb_00004auq, pdb_00004v3k, pdb_00004whv, pdb_00005bnb, pdb_00005d0k, pdb_00005d0m, pdb_00005dfl, pdb_00005eya, pdb_00005h7s, pdb_00005ifr, pdb_00005tut, pdb_00005ulf, pdb_00005ulh, pdb_00005ulk, pdb_00005vzw, pdb_00005zbu, pdb_00006jb6, pdb_00006jb7, pdb_00006s53, pdb_00006t7f, pdb_00006w9d, pdb_00007ai0, pdb_00007ai1, pdb_00007zj3, pdb_00008ams, pdb_00008eb0, pdb_00008grm, pdb_00008pjn, pdb_00008r5h, pdb_00008rx0, pdb_00009yea.

### Structure conformation and feature analysis

#### Closed/Open Conformation Classification

To classify ubiquitin–E2–E3 ternary complexes as either Closed or Open, we developed a workflow using the PyMOL python package (version 3.1.0) to determine the positioning of ubiquitin relative to the E2 protein. In particular, the distance between the Ile44 residue of ubiquitin and the E2 crossover helix was used to classify complexes as Closed or Open. For AF/CF predicted structures, which lack the anchoring ubiquitin-E2 conjugate bond, the distance between the ubiquitin tail and the E2 active site cysteine was also used to classify structures as Closed or Open. To identify the E2 crossover helix, a combination of secondary structure analysis and manual annotation was used to define helix start and stop residues. E2 active site cysteines were identified using PROSITE annotations and confirmed by manual review of E2 structures for positioning of any cysteines within the E2 after the UBC domain β-sheet and before the start of the crossover helix. For the Ile44-crossover helix distance, the minimum distance between the Cα atom of ubiquitin residue Ile44 and E2 crossover helix residue Cα atoms was calculated. For the ubiquitin tail to E2 cysteine distance, the distance from the carbonyl carbon of the ubiquitin C-terminal residue 76 and the Cα of the E2 active site cysteine was measured. Complexes within cutoffs of 10 Å for the Ile44–helix distance (and 12 Å for the ubiquitin C-terminus-cysteine for AF/CF modeled complexes) were designated Closed, all others as Open. For experimental structures in which the asymmetric unit contained multiple ternary complexes, distances were averaged across all ternary complexes in the asymmetric unit.

#### E2/E3 interaction and linchpin detection

RING/U-box-type E3s primarily bind with an E2 enzyme through interactions between loops within the E3 binding area of the E2 and loops of the E3 RING/U-box domain. As a prerequisite for assessing the complexes for E3 linchpin residues, we identified the E3 loops interacting with the E3-interacting loop of the E2. The E3-interacting loop of the E2 was identified as the coil region immediately N-terminal of the crossover helix. The E2 interaction region of the E3 was determined by including residues that fell within 8 Å of the E2 E3-binding loop. Secondary structure annotations, made using PyMOL define secondary structure (dss) function, were used to identify loop regions within the E3, and the number of interface residues that overlapped E3 loops was determined, as well as the number of E3 loops involved. Hydrogen bonding interactions between the C-terminal tail of ubiquitin (residues 71–76, or as much as this region was resolved in experimental structures), E2, and E3 components were assessed using distance-based detection of donor–acceptor atom pairs within 3.5 Å. PyMOL selections were used to extract relevant atomic coordinates, and all pairwise distances were computed to identify potential hydrogen bonds. All E3 residues within the E2-interacting region forming hydrogen bonds with the ubiquitin tail were identified and those residues that also formed hydrogen bonds with the E2 were flagged as linchpin residues. For experimental structures in which the asymmetric unit contained multiple ternary complexes, the structure was designated as linchpin-containing if at least half of the ternary complexes included such a positioned residue.

### Interface RMSD calculation

To quantify experimental and predicted ternary complex similarity, we calculated the root-mean-square deviation (RMSD) for interface residues. For experimental structures retrieved from the PDB, only the ubiquitin, E2, and E3 components of the ternary complex were considered, while any other proteins were excluded from the RMSD calculation. Predicted structures were generated using full-length ubiquitin and E2 enzyme sequences, while for E3_RING_ ligases, sequence input was limited to the RING domain-containing regions, and for RBR-type E3 ligases, the sequence corresponding to the region present in the experimental structure was used. Interface residues were defined as those with Cα atoms within 8.0 Å of a Cα atom from a different chain of the complex. Experimental and predicted structures were aligned for RMSD calculations using a Kabsch algorithm-based superposition implemented in the PDB module of the Biopython python package (version 1.85) via the Superimposer class, using residue pairs from the E2 crossover helices. All interface residues common to both the experimental and predicted structures were then used to calculate individual and overall interface RMSDs.

### STRING E2–E3 interactions

To identify E2–E3 interactions from the STRING database (88), we downloaded the protein network data for species *Homo sapiens* (STRING v12.0). The dataset was restricted to physical subnetwork only (i.e. inclusion of physical protein interactions only). We queried for any interactions of proteins in our list of E2s with proteins from our E3 list. For our pair-enriched E2–E3 set, only interactions with a STRING experimental evidence score of 400 (medium confidence according to STRING) or greater were included. For our non-pair enriched set, we combined all E2s and E3s from the paired set and removed all pairs with any level of STRING evidence for protein interaction. We restricted the non-pair set to only those E2s and E3s represented in the pair set to ensure that the non-pair set was not unfairly biased to poorly expressed or understudied proteins. For a final non-pair enriched set, a random selection was made to match the number of the pair enriched set.

### AlphaFold3 and ColabFold modeling

Modeling was performed on a high-performance computing cluster using NVIDIA A100 and NVIDIA L40S GPUs. Both AF3 (version 3.0.0) and ColabFold (CF) (version 1.5.5), were run in a containerized environment using Apptainer. A set of 50 models was generated for each complex using 10 different seeds (1-10 where specifiable), and 5 diffusion samples for each seed with AF3, and 5 model parameter sets in the case of CF. The prediction workflows were performed without templates, and both paired and unpaired multiple sequence alignments (MSAs) were utilized. Full ubiquitin and E2 sequences were used, while only RING/U-box domain containing truncations of E3 proteins were included. For RBR-type E3 RING proteins, the IBR portions were included (truncated sequences provided in ***SI Appendix,* Data S2**). For CF, a setting of 10 recycling iterations was applied with an early stopping tolerance of 0.5 (command line: colabfold_batch fasta_input output_folder--num-recycle=10--recycle-early-stop-tolerance=0.5--num-seeds=10). Both AF3 and CF models were relaxed with CF workflow using AMBER option and the first ranking model (per AF3/CF pipeline) was selected for subsequent analysis.

### Confidence score calculations

#### Interface Predicted Aligned Error (PAE)

For each protein complex, we calculated PAE values specific to interface regions. Interface residues were defined as those within a distance threshold of 8 Å from residue Cα atoms in an interacting chain, and a mean value was calculated. *Interface Predicted Score based on Aligned Errors (ipSAE)*: To further assess the predicted alignment error at protein–protein interfaces, we implemented the ipSAE metric which we previously developed for scoring protein-protein interactions of AlphaFold-based program predictions (77). In brief, for each pair of protein chains, all inter-chain residue pairs with a Predicted Alignment Error (PAE) below a defined threshold are included in the average of 1/(1+(PAE/d0)^2^), where *d*_0_ is calculated from the number of residues in the scored chain. ipSAE scores were specifically calculated using the publicly available Python script from the Dunbrack Lab (https://github.com/DunbrackLab/IPSAE). A PAE confidence threshold of 10 Å was used.

#### Predicted DockQ version 2 (pDockq2)

To quantify interface confidence based on predicted local distance difference test (pLDDT) and PAE scores, we implemented the pDockQ2 metric as described previously (78). pDockQ2 is a quantitative measure of interface quality calculated from integrating interface PAE and pLDDT values. Scores were specifically computed using the publicly available Python script from the Dunbrack Lab (https://github.com/DunbrackLab/IPSAE).

#### ipTM

For ipTM analysis, we used ipTM values for each of the three unique complex interfaces output by the AF3 workflow directly. For AlphaFold2 calculations, the pairwise ipTM values were obtained from the ipSAE code.

### Rosetta analysis

For calculation of interface binding energy, total and unsatisfied interface hydrogen bonds, total interface residues number, as well as total, polar, and non-polar SASA, we employed the InterfaceAnalyzer (79) application from the Rosetta software suite. InterfaceAnalyzer was run on relaxed structures using the default scoring function separately for each of the three interfaces individually, fixing two of the chains at a time.

### XGBoost classifier

#### Classifier training and evaluation

A supervised classification model using gradient-boosted decision trees implemented with XGBoost (88) was trained to classify E2–E3 combinations as pair or non-pair. Specifically, we used the XGBClassifier implementation from the XGBoost Python library (version 3.0.2). As input to the model, we included the results from our ternary complex analyses as features. This included confidence metrics (pairwise interface ipSAE, ipTM, pDockQ, pDockQ2 scores, as computed by the ipSAE Python analysis script), details of structural analysis (ubiquitin-Ile44 distance, ubiquitin-C-terminal tail distance, linchpin presence/absence, and number of E3 loops and residues interacting with E2) as well as the results of Rosetta InterfaceAnalyzer (pairwise interface binding ΔG, ΔSASA, hydrophobic/hydrophilic ΔSASA values, the number of total hydrogen bonds, unsatisfied hydrogen bonds, and total residues for each interface). Mean interface PAE values were excluded from the model input due to missing values for complexes lacking interface residues within the 8 Å cutoff used for their calculation, and redundancy to the other PAE-based metrics ipSAE and ipTM. Training-validation data complex labeling as ‘pair’ or ‘non-pair’ was based on pair vs non-pair sets derived from STRING data. To account for complexes removed from the initial pair and non-pair enriched sets, containing RBR-type E3s and non-functional E2s, we added additional randomly selected non-pair complexes so that the numbers of evaluated pair and non-pair complexes were equal. Complexes not in the Closed conformation were removed from the dataset before model training-validation. For feature filtering, we employed automated feature selection using SelectFromModel from scikit-learn (version 1.6.1) with the XGBoost classifier. Decision thresholds were selected by maximizing the F {0.75}-score. Performance was assessed using nested 5-fold stratified cross-validation. Metrics including accuracy, precision, recall, and area under the receiver operating characteristic curve (AUC-ROC) were computed to evaluate model performance. Due to the higher cross-validation performance of the CF-based model, we proceeded to train our final model for prediction on the complete CF dataset, using the optimal decision threshold determined from the cross-validation out-of-fold probabilities.

#### Prediction of E2 partners for E3 ligases

To predict potential E2 partners for E3_RING_ ligases lacking known E2 interactions, we selected all identified E3_RING_ proteins that had no interaction with any E2s in the STRING database (including all physical interactions of any evidence level). This resulted in a set of 88 E3_RING_ ligases. For each of these E3s, ternary complexes with ubiquitin and every E2 enzyme were modeled using CF as described above. Any structures in an Open conformation, lacking E2 engagement of the E3 RING/U-box domain, or with CF pTM or ipTM scores of less than 0.5 were filtered out and not included in subsequent analysis. All predicted structures were then processed using the same structural analysis pipeline as the training-validation dataset, including computation of model confidence scores, Rosetta InterfaceAnalyzer metrics, and additional structural features. These data were subsequently input into our CF-based trained model, to classify each modeled E2–E3 as either a predicted pair or non-pair.

### Statistics and reproducibility

Statistical analyses were performed using standard scientific Python libraries, including SciPy and NumPy. Specific statistical tests used are indicated in the figure legends and were chosen based on data structure and comparison type. All tests were two-sided unless otherwise noted. No statistical methods were used to predetermine sample size. No data were excluded from analyses, unless otherwise noted.

## Data Availability

The ColabFold-modeled ubiquitin–E2–E3 ternary complex structures generated in this study are available for download at our website, https://dunbrack.fccc.edu/ubiqcore.

## Acknowledgments

This work utilized computational resources from the Delta system at the National Center for Supercomputing Applications (NCSA) through allocation BIO250167 from the <u>Advanced</u> <u>Cyberinfrastructure Coordination Ecosystem: Services & Support</u> (ACCESS) (101) program, which is supported by U.S. National Science Foundation grants #2138259, #2138286, #2138307, #2137603, and #2138296. This work was also supported by NIH grant R35 GM122517 (R.L.D.).

## Author Contributions

R.L.D. and B.J. conceived the project. B.J. performed the analyses and prepared the figures. B.J. drafted the manuscript. R.L.D. reviewed and edited the manuscript.

## Supplementary Information Appendix

### Supplementary Tables

**Table S1.** Classification of proteins from culled E3 ligase lists. Atypical/Hybrid: E3 ligase *G2E3* has both HECT and RING domains

| **E3 Ligase Type** | **Count** | **Percentage (%)** |
| --- | --- | --- |
| **RING** | 318 | 54.83% |
| **HECT** | 28 | 4.83% |
| **RBR** | 14 | 2.41% |
| **U-box** | 8 | 1.38% |
| **Atypical/Hybrid** | 1 | 0.17% |
| **No E3 Catalytic Domain Detected** | 211 | 36.38% |

**Table S2.** Summary of experimental structure availability by E3 ligase type. E3 atypical: Atypical E3 ligase *G2E3* has both HECT and RING domains PDB query date: November 10, 2025

|  | With Structure | Without Structure | Total |
| --- | --- | --- | --- |
| E2 | 32 | 5 | 37 |
| E3 HECT | 21 | 7 | 28 |
| E3 RBR | 8 | 6 | 14 |
| E3 RING | 154 | 164 | 318 |
| E3 U-box | 6 | 2 | 8 |
| E3 atypical | 0 | 1 | 1 |
| Total | 221 | 185 | 406 |

**Table S3.** Summary of unique E2s and E3s in ternary complex experimental structures by type. Total ternary complex containing experimental structures, n=57 PDB query date: November 10, 2025

|  | In Ternary | Not in Ternary | Total |
| --- | --- | --- | --- |
| E2 | 11 | 26 | 37 |
| E3 HECT | 1 | 27 | 28 |
| E3 RBR | 5 | 9 | 14 |
| E3 RING | 23 | 295 | 318 |
| E3 U-box | 0 | 8 | 8 |
| E3 atypical | 0 | 1 | 1 |
| Total | 40 | 366 | 406 |

**Table S4.**
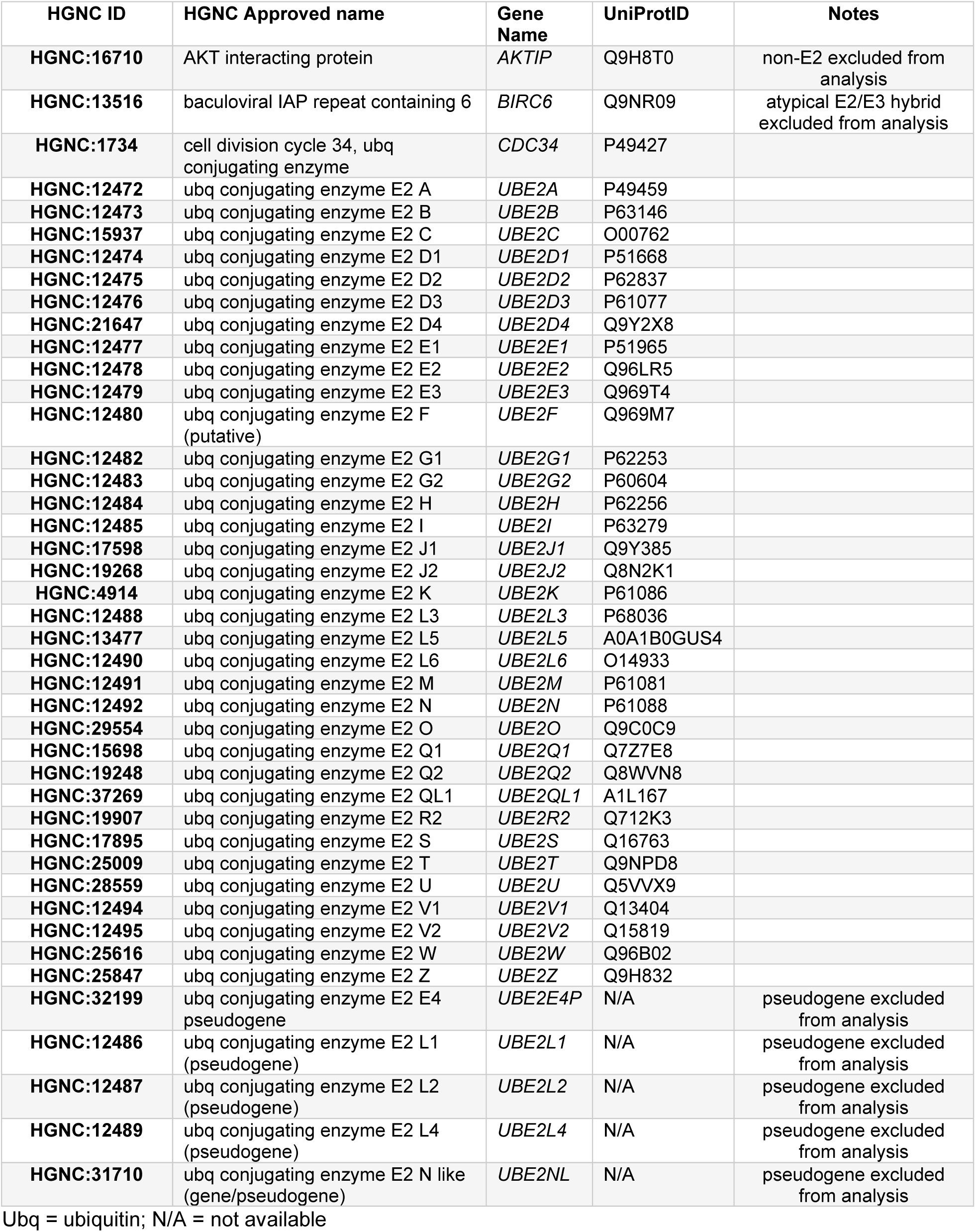
Retrieved E2 genes (HGNC Gene group: Ubiquitin conjugating enzymes E2 (UBE2)).

### Supplementary Figures

**Fig. S1:**
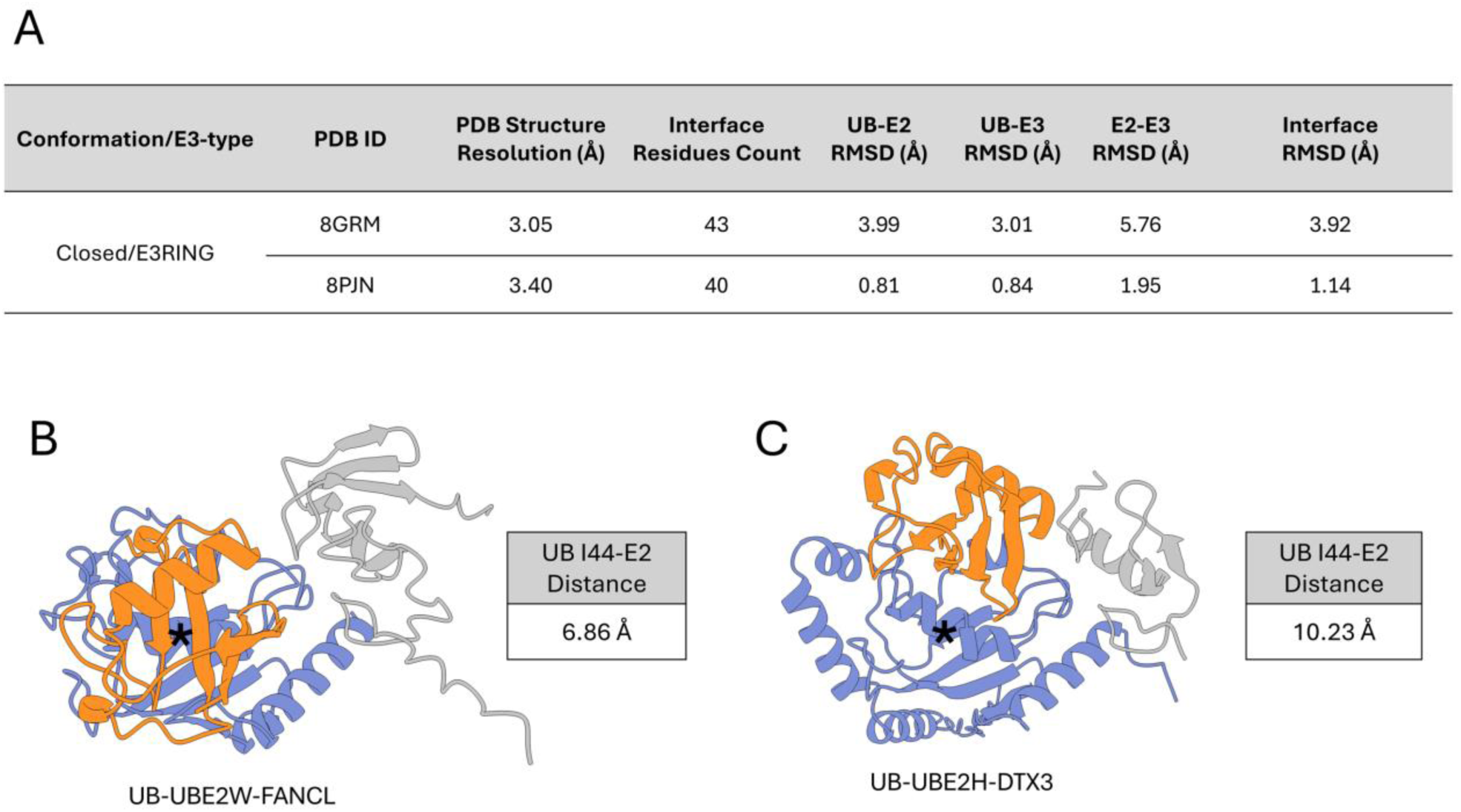
AF3 models of ubiquitin–E2–E3_RING_ ternary complexes without existing experimental structures with Closed/Open conformation assessment. (*A*) Table of interface RMSD values calculated by comparing experimentally determined structures of ubiquitin–E2–E3 complexes (PDB IDs listed) with their corresponding AF3-predicted models. In these AF3 predictions, the partner E3_RING_ of the heterodimers present in the experimental structures were explicitly included in the model alongside the core ubiquitin–E2–E3_RING_ components of the ternary complex. For 8GRM, the E3 RNF2, in a heterodimer with the E3 BMI1, makes primary contact with the E2 of the ternary complex (UBE2D2). For 8PJN, the E3 RMND5A, in a heterodimer with the E3 MAEA, makes primary contact with the E2 of the ternary complex (UBE2H). The table includes the resolution of the experimental structures, and the number of interface residues used in the RMSD calculations, defined as residues with Cα atoms within 8 Å across inter-protein interfaces of the complex. All experimental structures were released to the PDB after AF3 training dataset date cutoff of Sept. 30, 2021 (*B*) AF3 predicted structure of ubiquitin in complex with E2, UBE2W and E3, FANCL, a putative functional E2–E3 pair. This ternary complex is in the Closed conformation, based on determined 10 Å Ub Ile44-crossover helix distance cutoff. (*C*) AF3 predicted structure of ubiquitin in complex with E2, UBE2H and E3, DTX3, an E2 and E3 with no evidence of interaction. This ternary complex is in an Open conformation, based on determined 10 Å Ub Ile44-crossover helix distance cutoff. Structures were retrieved from the RCSB Protein Data Bank and visualized using UCSF ChimeraX version 1.10.1. Asterisk (*) indicates crossover helix of the E2 in the ternary complexes.

**Fig. S2:**
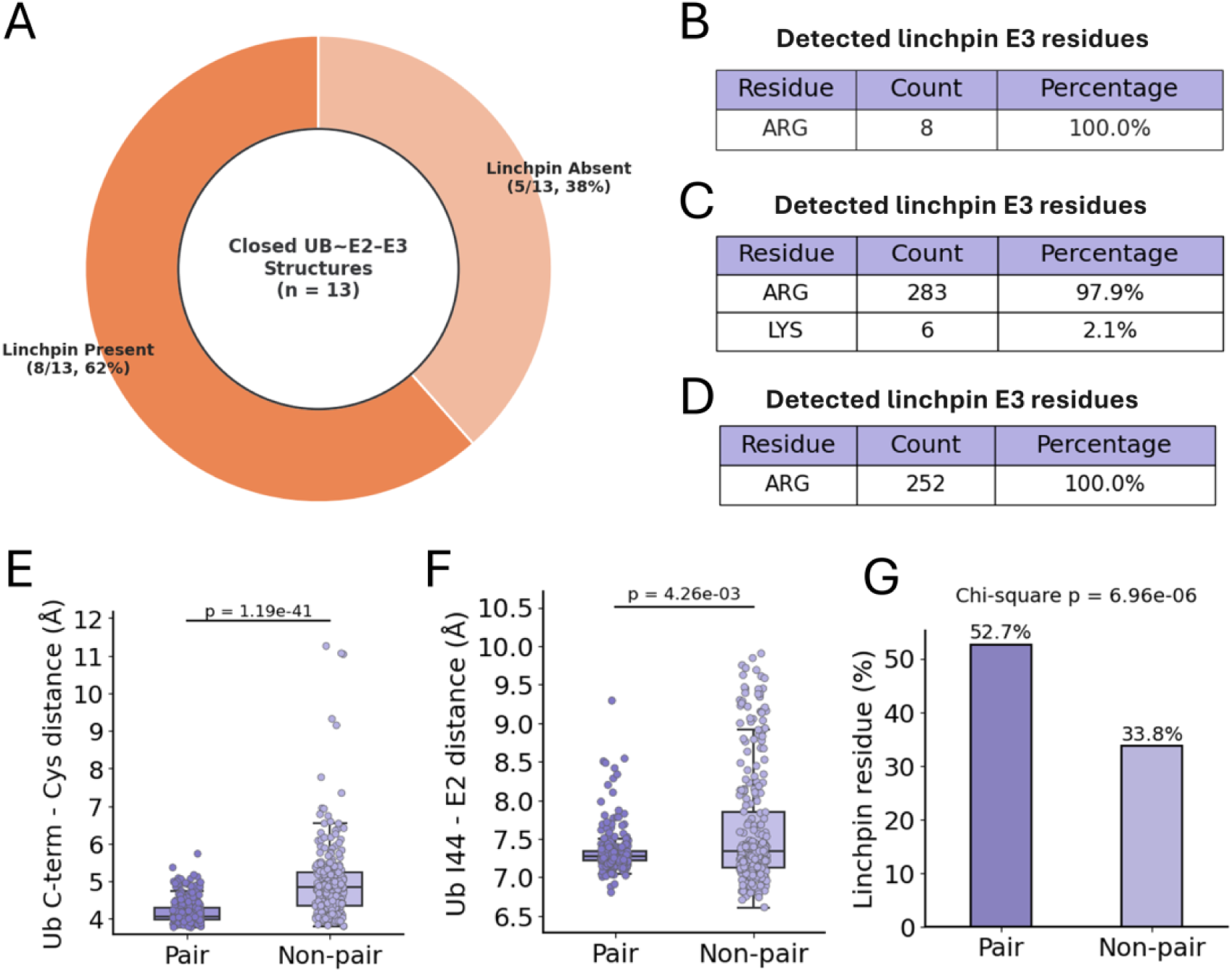
Analysis of linchpin residues in experimental, AF3, and CF models ternary complexes, and additional structural features of pair and non-pair ubiquitin–E2–E3 Closed conformation ternary complex structures generated with ColabFold. (*A*) Proportion of Closed conformation experimental ubiquitin∼E2–E3_RING_ ternary complexes featuring an E3 ligase linchpin residue. (*B*) Table of detected E3 linchpin residue types in Closed conformation experimental ubiquitin∼E2–E3_RING_ ternary complexes. (*C*) Table of detected E3 linchpin residue types across AF3 modeled ternary complexes of both pair and non-pair E2–E3 sets. (*D*) Table of detected E3 linchpin residue types across CF modeled ternary complexes of both pair and non-pair E2–E3 sets. (*E)* Distances from ubiquitin Ile44 Cα to closest crossover helix Cα in CF modeled ternary complexes, p-value calculated using Mann-Whitney U Test (two-sided). (*F*) Distances from ubiquitin terminal glycine Cα to the E2 active-site cysteine carbonyl carbon Cα in CF modeled ternary complexes, p-value calculated using Mann-Whitney U Test (two-sided). (*G*) Proportion of CF modeled ternary complexes featuring linchpin residue in pair and non-pair complexes. (*E-G*) *n* = 298 (pair) and *n* = 281 (non-pair).

**Fig. S3:**
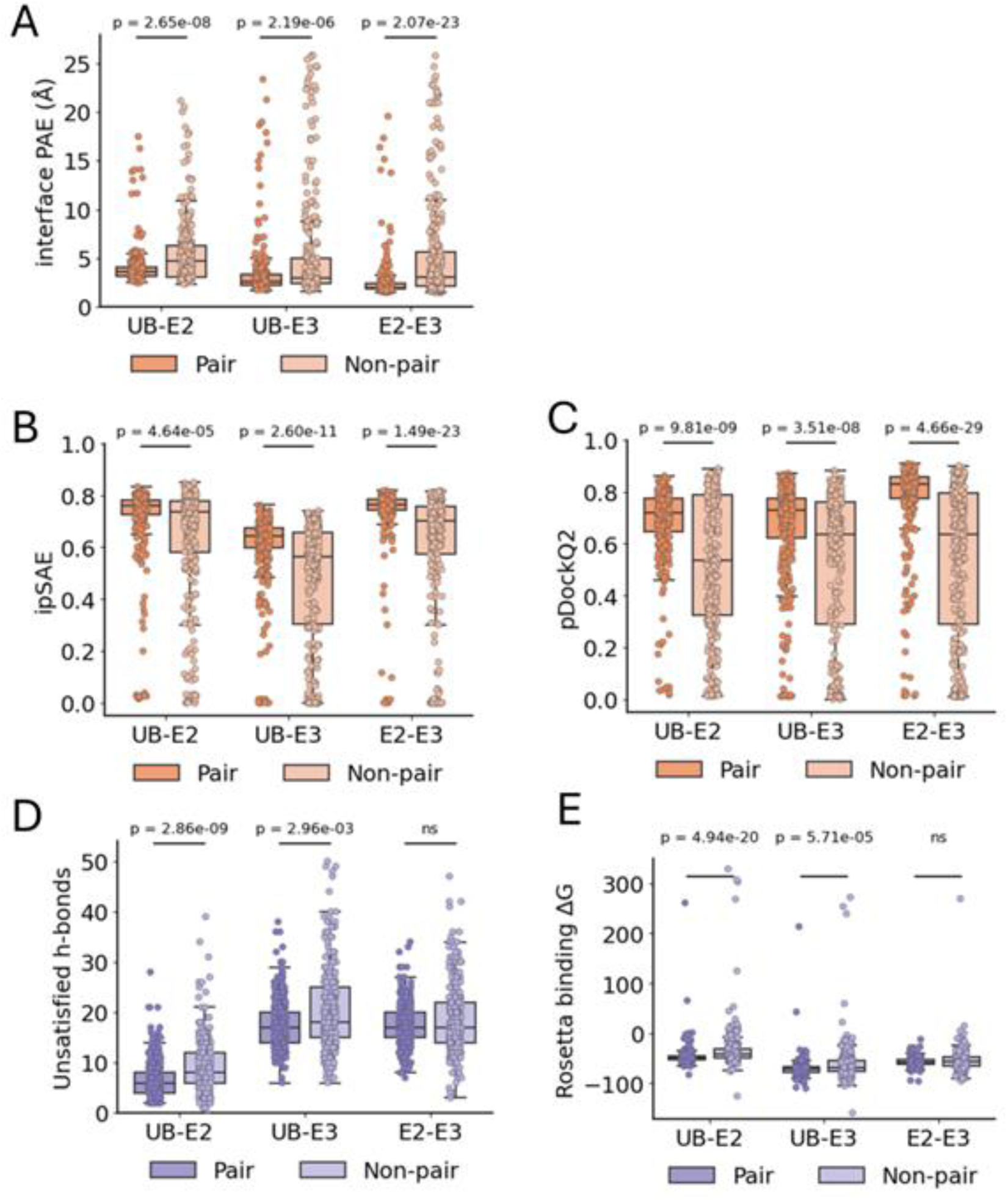
ColabFold modeling of ubiquitin–E2–E3 ternary complexes of E2–E3 pair and non-pair sets with Closed/Open conformation analysis. (*A*) Scatter plot of ubiquitin Ile44 to E2 crossover helix distance vs ubiquitin Gly C-terminus to E2 catalytic cysteine distance for all structures. (Note: Structures containing the E2s UBE2V1 or UBE2V2, which lack a catalytic cysteine, are not shown in this plot. However, all such structures exceeded the 10 Å I44–E2 crossover helix distance cutoff and were assigned Open conformation designation.) (*B*) Proportion of ternary complexes in a Closed vs. Open conformation for pair and non-pair set. Closed/Open proportion by E2 (*E*), E3_RBR_ (RBR-type E3) (*C*), and the 30 lowest Closed proportion ranking E3_RING_ (RING/U-box-type E3) (*D*) (E2s and E3s referred to by gene name).

**Fig. S4:**
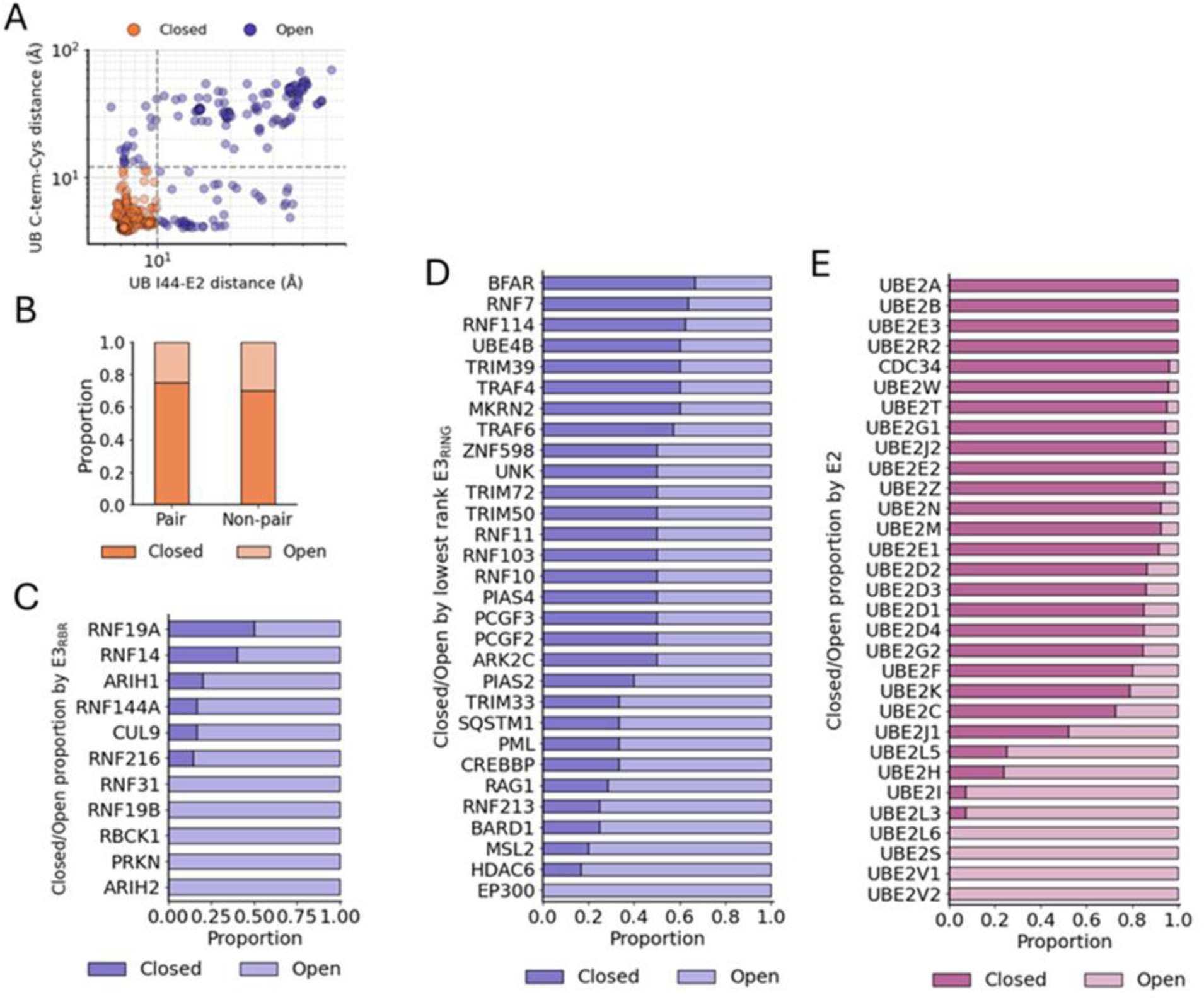
Interface confidence metrics, energetics, and interactions of pair and non-pair ubiquitin–E2–E3 Closed conformation ternary complex structures generated with ColabFold. Average interface PAE values (based on 8 Å Cα-Cα interface cutoff) (*A*), ipSAE (*B*), and pDockq2 (*C*) plotted for each complex interface. Number of unsatisfied hydrogen bonds at interfaces (*D*) and complex interface binding energies (Rosetta units) (*E*), both determined using Rosetta InterfaceAnalyzer. Two-sided Mann-Whitney U tests were used to calculate p-values; ns denotes p > 0.05. For panels b–e, *n* = 298 (pair) and *n* = 281 (non-pair). For panel a, group sizes vary by interface (due to exclusion of structures lacking interface residues within the 8 Å cutoff): UB–E2: *n* = 298 (pair), *n* = 281 (non-pair); UB–E3: *n* = 298 (pair), *n* = 275 (non-pair); E2–E3: *n* = 298 (pair), *n* = 281 for non-pair.

